# Identification of signaling pathways, matrix-digestion enzymes, and motility components controlling *Vibrio cholerae* biofilm dispersal

**DOI:** 10.1101/2020.10.09.333351

**Authors:** Andrew A. Bridges, Chenyi Fei, Bonnie L. Bassler

## Abstract

Bacteria alternate between being free-swimming and existing as members of sessile multicellular communities called biofilms. The biofilm lifecycle occurs in three stages: cell attachment, biofilm maturation, and biofilm dispersal. *Vibrio cholerae* biofilms are hyper-infectious and biofilm formation and dispersal are considered central to disease transmission. While biofilm formation is well-studied, almost nothing is known about biofilm dispersal. Here, we conduct an imaging screen for *V. cholerae* mutants that fail to disperse, revealing three classes of dispersal components: signal transduction proteins, matrix-degradation enzymes, and motility factors. Signaling proteins dominated the screen and among them, we focused on an uncharacterized two-component sensory system that we name DbfS/DbfR for Dispersal of Biofilm Sensor/Regulator. Phospho-DbfR represses biofilm dispersal. DbfS dephosphorylates and thereby inactivates DbfR, which permits dispersal. Matrix degradation requires two enzymes: LapG, which cleaves adhesins, and RbmB, which digests matrix polysaccharide. Reorientations in swimming direction, mediated by CheY3, are necessary for cells to escape from the porous biofilm matrix. We suggest that these components act sequentially: signaling launches dispersal by terminating matrix production and triggering matrix digestion and, subsequently, cell motility permits escape from biofilms. This study lays the groundwork for interventions that modulate *V. cholerae* biofilm dispersal to ameliorate disease.

**Significance statement:** The pathogen *Vibrio cholerae* alternates between the free-swimming state and existing in sessile multicellular communities known as biofilms. Transitioning between these lifestyles is key for disease transmission. *V. cholerae* biofilm formation is well studied, however, almost nothing is known about how *V. cholerae* cells disperse from biofilms, precluding understanding of a central pathogenicity step. Here, we conducted a high-content imaging screen for *V. cholerae* mutants that failed to disperse. Our screen revealed three classes of components required for dispersal: signal transduction, matrix degradation, and motility factors. We characterized these components to reveal the sequence of molecular events that choreograph *V. cholerae* biofilm dispersal. Our report provides a framework for developing strategies to modulate biofilm dispersal to prevent or treat disease.

## Main

Bacteria transition between existing in the biofilm state, in which cells are members of surface-associated multicellular collectives, and living as free-swimming, exploratory individuals. Biofilms consist of cells surrounded by a self-secreted extracellular matrix that protects the resident cells from threats including predation, antimicrobials, and dislocation due to flow.(1–3) Biofilms are relevant to human health because beneficial microbiome bacteria exist in biofilms, and, during disease, because pathogens in biofilms evade host immune defenses, thwart medical intervention, and exhibit virulence.(4–7) The biofilm lifecycle consists of three stages: cell attachment, biofilm maturation, and dispersal (Figure 1A).(8) Cells liberated during the dispersal step can disseminate and found new biofilms.(8) The environmental stimuli and the components facilitating biofilm attachment and maturation have been defined for many bacterial species.(9) In contrast, little is known about the biofilm dispersal stage.

**Figure 1.**
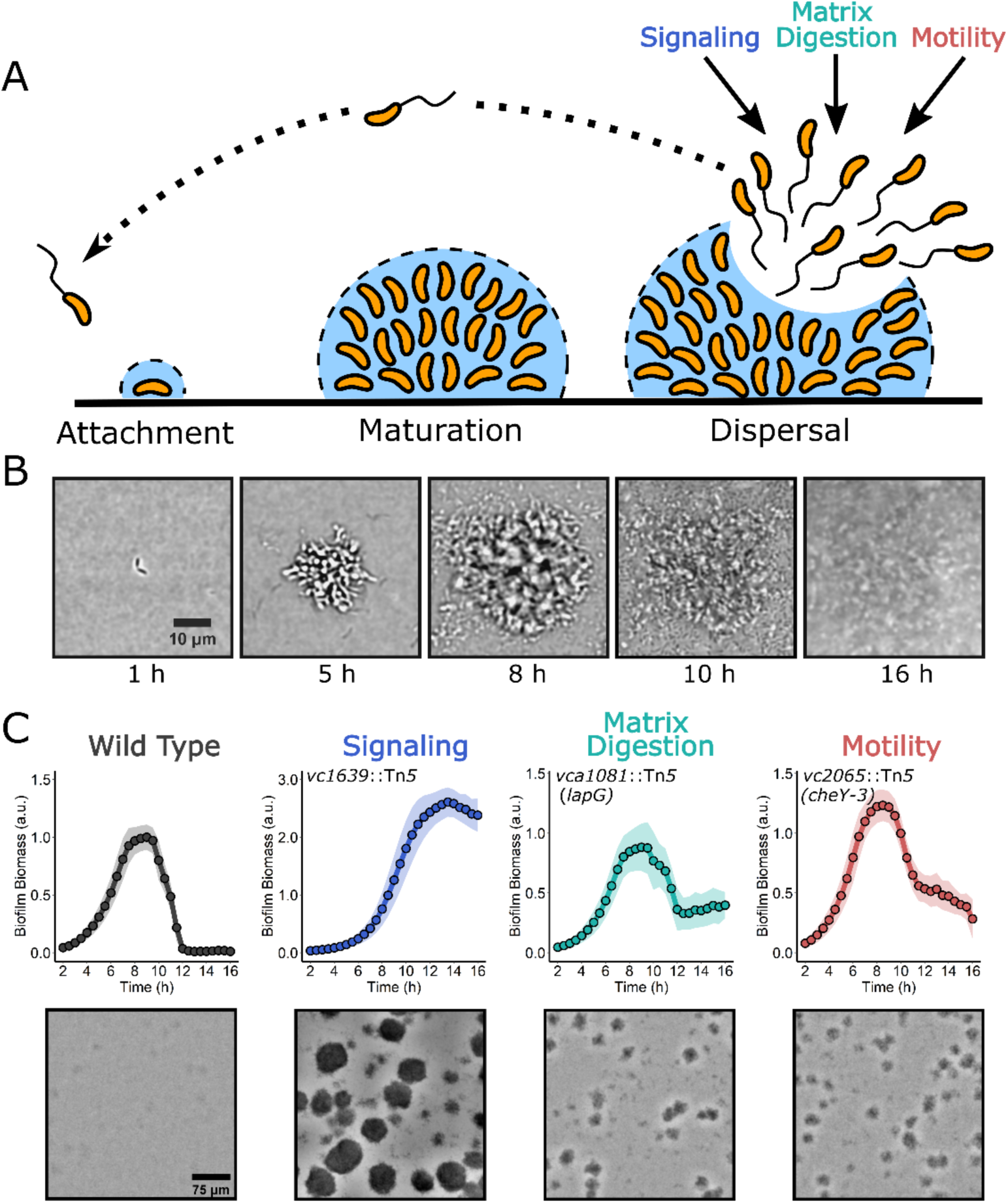
A high-content imaging screen identifies genes required for *V. cholerae* biofilm dispersal. (A) Schematic illustrating the *V. cholerae* biofilm lifecycle. See text for details. (B) Brightfield image series over time of the WT *V. cholerae* biofilm lifecycle. (C) Top panels: Quantitation of biofilm biomass over time as measured by time-lapse microscopy for WT and representative transposon insertion mutants from each of the three functional categories identified in the screen. Note differences in *y*-axes scales. Data are represented as means normalized to the peak biofilm biomass of the WT strain. *N* = 3 biological and *N* = 3 technical replicates, ± SD (shaded). a.u., arbitrary unit. Bottom panels: Representative brightfield images of biofilms at the final 16 h timepoint for the strains presented in the top panels.

The model pathogen *Vibrio cholerae* forms biofilms in its aquatic habitat, biofilm cells are especially virulent in mouse models of cholera disease, and biofilms are thought to be critical for cholera transmission.(10–14) Studies of *V. cholerae* biofilms have predominantly focused on matrix overproducing strains that constitutively exist in the biofilm mode and that do not disperse. This research strategy has propelled understanding of *V. cholerae* biofilm attachment and maturation, revealing that the second messenger cyclic diguanylate (c-di-GMP) is a master regulator of biofilm formation, and that expression of vibrio polysaccharide (*vps*) biosynthetic genes are required.(15–17) The strategy of characterizing constitutive biofilm formers, while successful for uncovering factors that promote biofilm formation, has necessarily precluded studies of biofilm dispersal. Here, we employed a microscopy assay that allowed us to monitor the full wild-type (WT) *V. cholerae* biofilm lifecycle. We combined this assay with high-content imaging of randomly mutagenized WT *V. cholerae* to identify genes required for biofilm dispersal. Investigation of the proteins encoded by the genes allowed us to characterize the signaling relays, matrix-digestion enzymes, and motility components required for biofilm dispersal, a key stage in the lifecycle of the global pathogen *V. cholerae*.

## Results

Previously, we developed a brightfield microscopy assay that allows us to monitor the full WT *V. cholerae* biofilm lifecycle in real time.(18) In our approach, *V. cholerae* cells are inoculated onto glass coverslips at low cell density and brightfield time-lapse microscopy is used to monitor biofilm progression. WT biofilms reach peak biomass after 8-9 h of incubation and subsequently dispersal occurs and is completed by 12-13 h (Figure 1B, C). To identify genes required for biofilm dispersal, we combined mutagenesis with high-content imaging of the output of this assay. Specifically, WT *V. cholerae* was mutagenized with Tn*5* yielding ∼7000 mutants that were arrayed in 96-well plates. Following overnight growth, the mutants were diluted to low cell density in minimal medium, a condition that drives initiation of the biofilm lifecycle. Brightfield images of each well were captured 8 h post-inoculation to assess biofilm maturation and at 13 h to evaluate biofilm dispersal. Mutants that showed no defects in biofilm maturation as judged by the 8 h images but displayed significant remaining biofilm biomass at the 13 h timepoint were identified. To verify phenotypes, candidate mutants were individually reevaluated by time-lapse microscopy. Mutants that accumulated at the bottom of wells due to aggregation or that failed to attach to surfaces were excluded from further analysis, eliminating strains harboring insertions in O-antigen and flagellar genes, respectively. The locations of transposon insertions in the 47 mutants that met our criteria were defined and corresponded to 10 loci. The new genes from the screen fell into three classes: signal transduction (blue), matrix degradation (green), and motility (red) (Figure 1A, C). In-frame deletions of each gene were constructed, and the biofilm lifecycles of the deletion mutants were imaged to confirm that the genes are required for biofilm dispersal (Table 1, Video 1). We also identified insertions in genes encoding proteins with known roles in biofilm dispersal (i.e., RpoS, quorum sensing), which we excluded from further analysis.(18, 19)

**Table 1:**
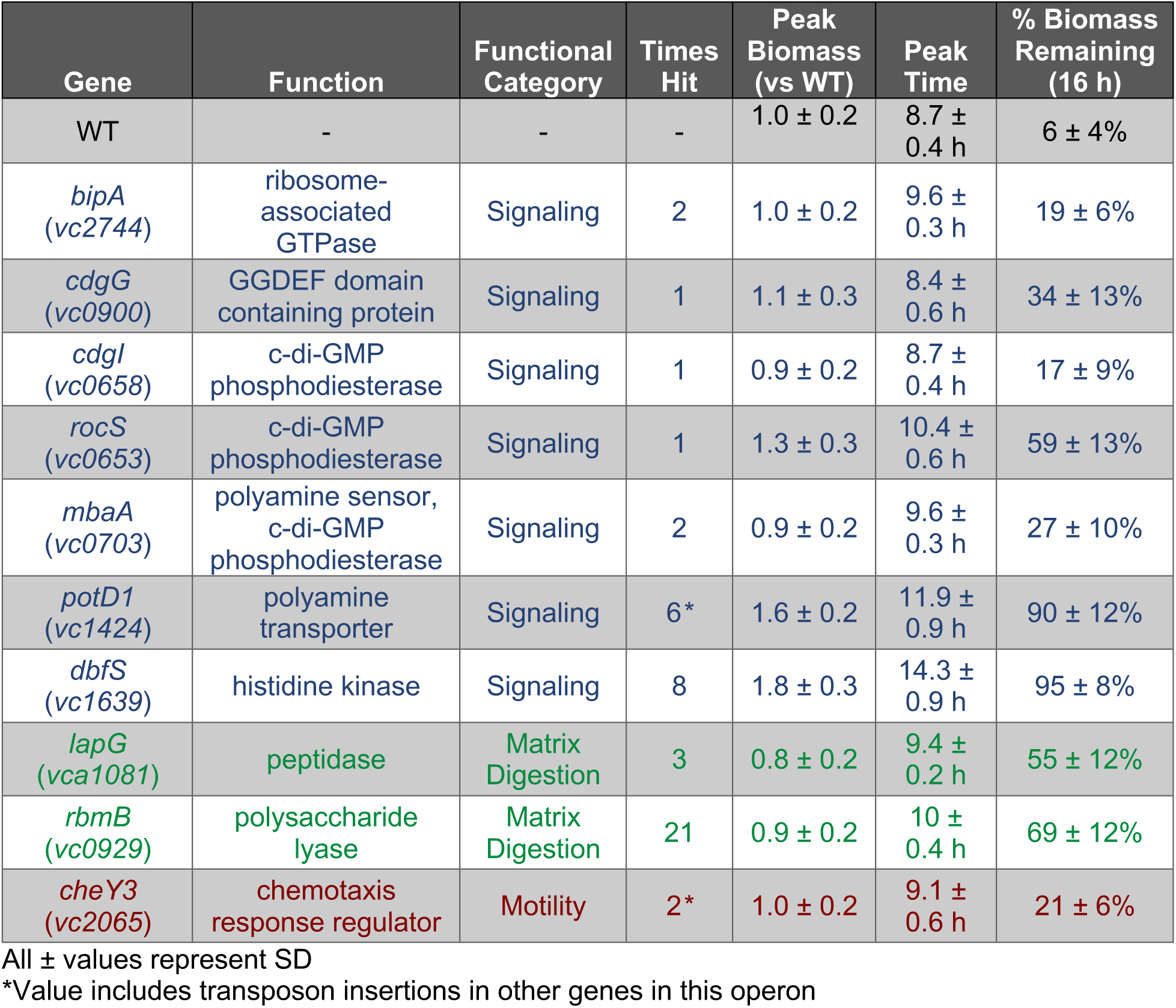
Genes identified as required for *V. cholerae* biofilm dispersal and phenotypes of deletion mutants.

Proteins involved in signal transduction dominated the screen (7 of 10 loci) and included the ribosome-associated GTPase, BipA, multiple cyclic diguanylate (c-di-GMP) signaling proteins, polyamine signaling proteins, and a putative two-component histidine kinase, Vc1639. The signal transduction mutants displayed different severities in their biofilm dispersal phenotypes. The *ΔbipA* displayed a modest defect: ∼19% of its biofilm biomass remained at 16 h, the final timepoint of our data acquisition, while the WT showed ∼6% biomass remaining. By contrast, the *Δvc1639* mutant underwent no appreciable dispersal (Table 1). In the category of matrix degradation, two enzymes were identified, LapG a periplasmic protease, and RbmB, a putative polysaccharide lyase (Table 1). A single motility mutant was identified with an insertion in the gene encoding the chemotaxis response regulator *cheY3* (Table 1). Below, we carry out mechanistic studies on select mutants from each category to define the functions of the components. Other mutants will be characterized in separate reports.

### A two-component regulatory system controls *V. cholerae* biofilm dispersal

The mutant from our screen that exhibited the most extreme dispersal phenotype had a transposon in a gene encoding an uncharacterized putative histidine kinase (designated HK), Vc1639 (Table 1). A screen for factors required for *V. cholerae* colonization of the suckling mouse intestine repeatedly identified Vc1639, suggesting that this HK is core to the cholera disease.(20) HKs typically contain periplasmic ligand binding domains and internal catalytic domains that switch between kinase and phosphatase activities based on ligand detection.(21) HKs transmit sensory information to cognate response regulators (RR) by altering RR phosphorylation.(22) RRs, in turn, control gene expression and/or behavior depending on their phosphorylation states. Deletion of *vc1639* in *V. cholerae* resulted in an 80% increase in peak biofilm biomass relative to WT and nearly all the biofilm biomass remained at 16 h demonstrating that Vc1639 is essential for biofilm dispersal (Figure 2A, Table 1). Complementation of the Δ*vc1639* mutant with *vc1639* inserted onto the chromosome at an ectopic locus restored WT biofilm dispersal (Supplementary Figure 1A). Consistent with the extreme dispersal phenotype of the Δ*vc1639* mutant, *vpsL-lux* expression was elevated 10-fold throughout the growth curve in the Δ*vc1639* strain compared to WT *V. cholerae* (Figure 2B). *vpsL* is the first gene in the major extracellular matrix biosynthetic operon showing that Vc1639 signaling regulates matrix production. Likewise, *lux* promoter fusions to the genes encoding the biofilm master regulators *vpsR* and *vpsT* also exhibited increased light production in the Δ*vc1639* mutant suggesting that VC1639 acts at the top of the cascade to control global biofilm gene expression (Supplementary Figure 1B, C). *vc1639* is the final gene in a three gene operon that includes genes encoding a hypothetical protein (Vc1637) and an OmpR family RR (Vc1638) (Figure 2C). We name Vc1639 DbfS for Dispersal of Biofilm Sensor and we name Vc1638 DbfR for Dispersal of Biofilm Regulator. Domain prediction suggests that DbfS contains two transmembrane domains (TM), a periplasmic sensory domain, and a cytoplasmic HAMP domain that likely transmits ligand-binding-induced conformational changes to regulation of the C-terminal kinase/phosphatase activity (Figure 2C).

**Figure 2.**
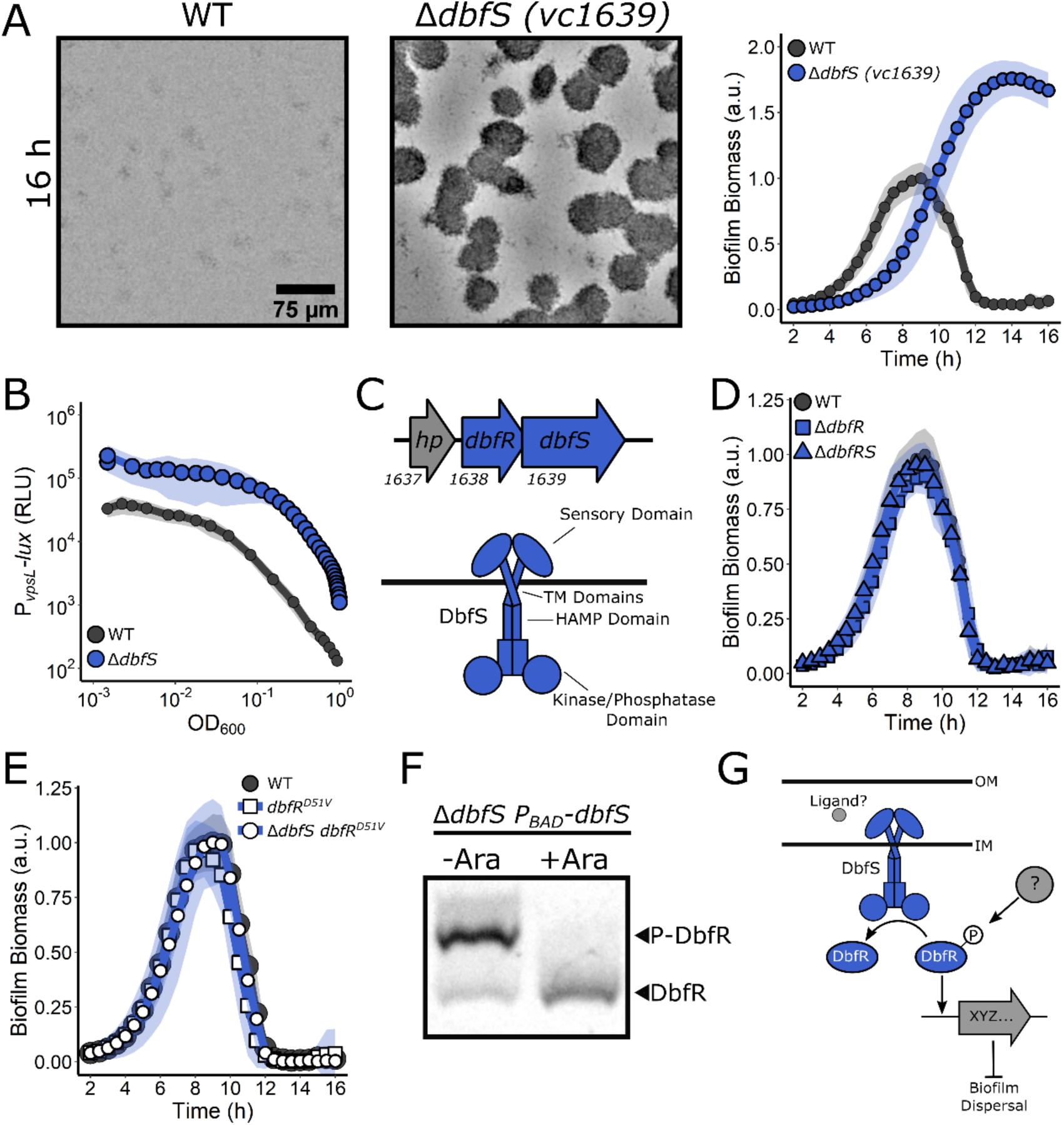
A two-component system composed of DbfS (HK) and DbfR (RR) controls *V. cholerae* biofilm dispersal. (A) Representative 16 h images and quantitation of biofilm biomass over time measured by time-lapse microscopy for WT *V. cholerae* and the Δ*dbfS* (i.e., Δ*vc1639*) mutant. (B) The corresponding *P*_*vpsL*_*-lux* output for strains and growth conditions in A over the growth curve. (C) Top panel: operon structure of the genes encoding the DbfS-DbfR two-component system. Bottom panel: Cartoon of the domain organization of DbfS. TM, transmembrane domain (D) As in A for the Δ*dbfR* (i.e., Δ*vc1638*) strain and for the Δ*dbfS* Δ*dbfR* double mutant. (E) As in A for the *dbfR*^*D51V*^ and Δ*dbfS dbfR*^*D51V*^ strains. (F) Representative Phos-tag gel analysis of DbfR-SNAP in the absence (-arabinose) or presence (+arabinose) of DbfS. Fucose was added to repress DbfR production in the uninduced samples. A phosphorylated protein migrates slower than the same unphosphorylated protein. (G) Proposed model for the DbfS-DbfR phosphorylation cascade regulating biofilm dispersal. OM, outer membrane; IM, inner membrane. In all biofilm measurements, *N* = 3 biological and *N* = 3 technical replicates, ± SD (shaded). a.u., arbitrary unit. For *vpsL-lux* measurements, *N* = 3 biological replicates, ± SD (shaded). RLU, relative light units. Phos-tag gel result is representative of *N* = 3 independent biological replicates.

To explore the connection between DbfS and DbfR in the control of biofilm dispersal, we deleted *dbfR*. Commonly, cognate HK and RR null mutants have identical phenotypes. To our surprise, the Δ*dbfR* mutant had no biofilm dispersal defect and progressed through the biofilm lifecycle identically to WT (Figure 2D). We considered the possibility that some other RR is the partner to DbfS. To test this idea, we constructed the Δ*dbfS* Δ*dbfR* double mutant. This strain behaved identically to the Δ*dbfR* strain (Figure 2D), demonstrating that *dbfR* is epistatic to *dbfS* and thus, DbfR indeed functions downstream of DbfS. Moreover, because RRs are typically active when phosphorylated, our results suggest that DbfR must be active in the absence of DbfS. Thus, we reason that phospho-DbfR is the species present in the Δ*dbfS* strain. To verify the hypothesis that phospho-DbfR is responsible for the dispersal defect in the Δ*dbfS* strain, we constructed a non-phosphorylatable allele of DbfR (D51V). The *V. cholerae dbfR*^*D51V*^ mutant displayed the WT biofilm dispersal phenotype in the presence and the absence of DbfS (Figure 2E). DbfR-SNAP fusions showed that SNAP did not interfere with WT DbfR function and that DbfR protein abundance was unchanged in the *dbfR*^*D51V*^ strain relative to WT (Supplementary Figure 1D, E). Thus, phospho-DbfR causes *V. cholerae* cells to remain in the biofilm state in the Δ*dbfS* mutant. It follows that deletion of *dbfS* causes biofilm dispersal failure due to loss of DbfS phosphatase activity on DbfR. To test this hypothesis, we assessed *in vivo* DbfR phosphorylation in the presence and absence of DbfS. Phos-tag gel analysis enabled separation and visualization of phosphorylated and dephosphorylated DbfR. In the absence of DbfS, DbfR was phosphorylated and induction of DbfS production caused the phospho-DbfR species to disappear (Figure 2F). Thus, under our experimental conditions, DbfS functions as a DbfR phosphatase. We infer that some other unknown kinase must exist and phosphorylate DbfR (Figure 2G). We propose that phospho-DbfR is active, and it drives expression of matrix biosynthetic genes, and increased matrix production prevents biofilm dispersal. It is possible that phospho-DbfR also controls other genes involved in suppressing biofilm dispersal.

DbfS is well-conserved in the vibrio genus, for example, in *Vibrio vulnificus* and *Vibrio parahaemolyticus*, DbfS has respectively, 64% and 60% amino acid sequence identity to *V. cholerae* DbfS. In genera closely related to vibrio, i.e., allovibrio and photobacteria, the *dbfS* gene exists in an identical operon organization and the encoded protein shows high amino acid sequence identity (∼55-65%) to *V. cholerae* DbfS. In many cases, *dbfS* is annotated as *phoQ*, encoding the well-studied cation-regulated HK from enteric pathogens including *Escherichia coli* and *Salmonella*. However, BLAST analysis of the DbfS protein sequence against that from *E. coli* K-12 revealed limited homology to PhoQ, with 32% amino acid sequence identity (E value=1e^-41^), with the lowest region of similarity in the predicted ligand binding domain. We tested whether the ligands that control PhoQ signal transduction also regulate DbfS-DbfR signaling (Supplementary Figure 2A-D, Supplemental Discussion). They do not. Thus, DbfS and DbfR are not functionally equivalent to PhoQ and its cognate RR, PhoP, respectively. Thus, DbfS responds to a yet-to-be defined stimulus to regulate biofilm dispersal.

### Matrix disassembly mediates *V. cholerae* exit from biofilms

The second group of mutants in our screen harbored insertions in the gene encoding the calcium-dependent periplasmic protease LapG that degrades outer-membrane spanning adhesive proteins and in the gene specifying the extracellular polysaccharide lyase RbmB that degrades the VPS component of the biofilm matrix.(23, 24) The Δ*lapG* strain exhibited slightly lower peak biofilm biomass compared to WT, with a short delay in the onset of dispersal, and ∼55% of its biomass remained at 16 h (Figure 3A, Table 1). The Δ*lapG* and the WT strains had similar *vpsL-lux* expression patterns (Figure 3B) consistent with LapG playing no role in repression of matrix production, but rather functioning downstream in matrix degradation. The LapG mechanism is known: When c-di-GMP concentrations are high, the FrhA and CraA adhesins are localized to the outer membrane where they facilitate attachments that are important for biofilm formation (Figure 3C).(25, 26) Under this condition, LapG is sequestered and inactivated by the inner membrane c-di-GMP sensing protein LapD.(25) When c-di-GMP levels fall, LapD releases LapG, and LapG cleaves FrhA and CraA facilitating cell detachment from biofilms.(25) Our results are consistent with this mechanism; in the absence of LapG, FrhA and CraA remain intact, and *V. cholerae* cells cannot properly exit the biofilm state. To verify that the established c-di-GMP-dependent regulatory mechanism controls LapG activity in our assay, we deleted *lapD* (Figure 3C). Indeed, in the Δ*lapD* strain, biofilm dispersal occurred prematurely indicating that, without LapD, LapG is not sequestered, and unchecked LapG activity promotes premature adhesin degradation, and, as a consequence, early biofilm disassembly (Figure 3D). The Δ*lapD* Δ*lapG* double mutant had the same dispersal phenotype as the Δ*lapG* single mutant confirming that LapG functions downstream of LapD (Figure 3D). Lastly, in a reciprocal arrangement, overexpression of *lapG* from an ectopic locus caused peak biofilm formation to decrease by ∼65% (Supplementary Figure 3A) suggesting that enhanced LapG-mediated cleavage of adhesins prematurely released cells from the biofilm. Thus, the conserved Lap pathway, which responds to changes in c-di-GMP levels, facilitates biofilm dispersal in *V. cholerae*.

**Figure 3.**
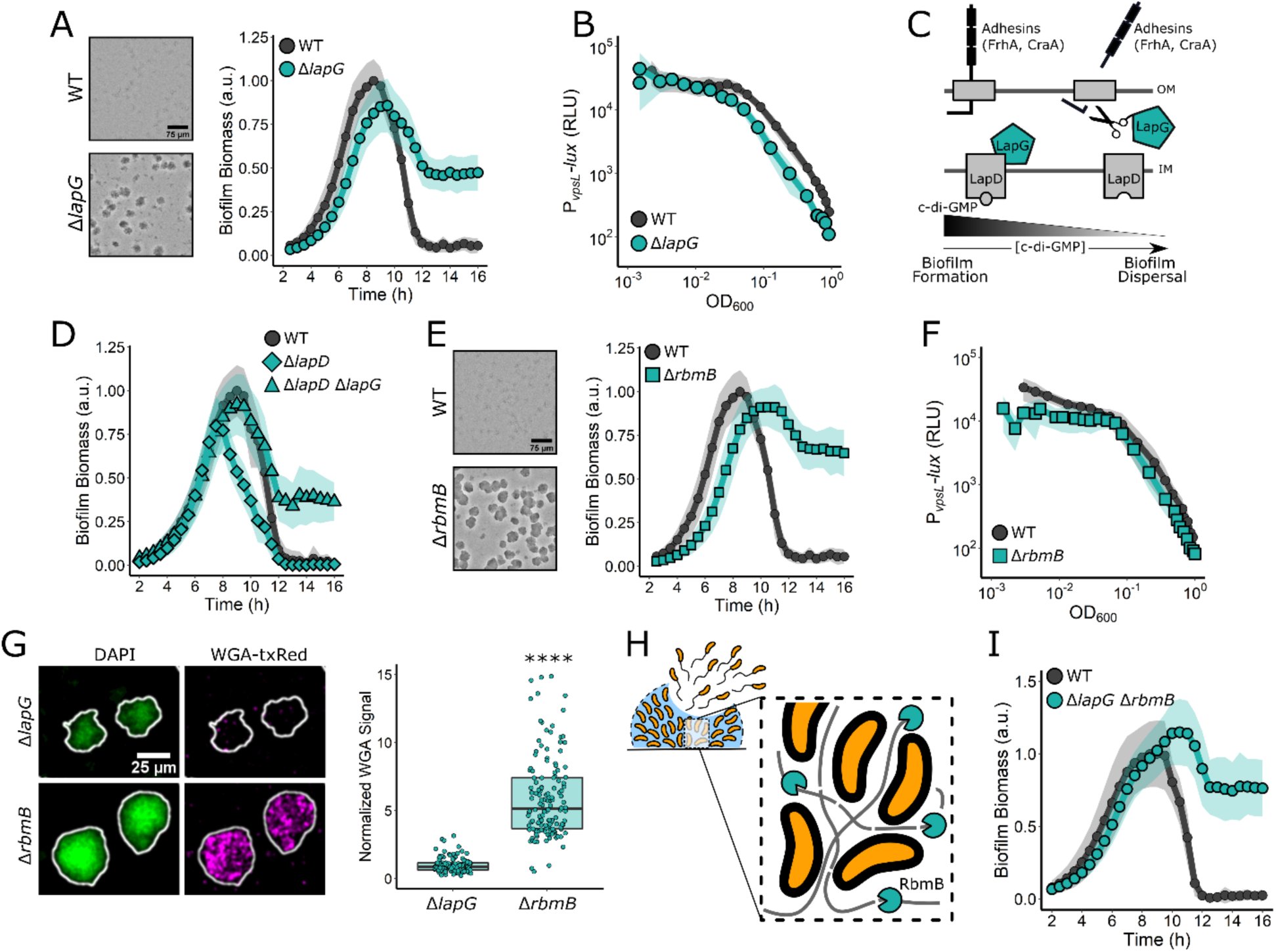
Matrix-digesting enzymes mediate *V. cholerae* biofilm dispersal. (A) Representative 16 h images and quantitation of biofilm biomass over time measured by time-lapse microscopy for WT *V. cholerae* and the Δ*lapG* mutant. (B) The corresponding *P*_*vpsL*_*-lux* output for strains and growth conditions in A over the growth curve. (C) Schematic representing the LapG mechanism. (D) As in A for the WT, the Δ*lapD* single mutant, and the Δ*lapD* Δ*lapG* double mutant. (E) As in A for the WT and the Δ*rbmB* mutant. (F) As in B for WT *V. cholerae* and the Δ*rbmB* mutant. (G) Representative images and quantitation of WGA-txRed signal in Δ*lapG* and Δ*rbmB* biofilms 16 h post-inoculation. To account for differences in biomass, the WGA-txRed signal was divided by the 4’, 6-diamidino-2-phenylindole (DAPI) signal in each biofilm. Values were normalized to the mean signal for the Δ*lapG* strain. >100 individual biofilms were quantified for each strain. An unpaired t-test was performed for statistical analysis, with **** denoting p < 0.0001. (H) Proposed model for the role of RbmB in biofilm dispersal. Gray lines represent the polysaccharide matrix. (I) As in A for the WT and the Δ*lapG* Δ*rbmB* double mutant. In all cases, *N* = 3 biological and *N* = 3 technical replicates, ± SD (shaded). a.u., arbitrary unit. For *vpsL-lux* measurements, *N* = 3 biological replicates, ± SD (shaded). RLU, relative light units. OM, outer membrane; IM, inner membrane.

Regarding the RbmB polysaccharide lyase, the Δ*rbmB* strain formed biofilms to roughly the same peak biomass as WT, however, it exhibited a 2 h delay in dispersal onset and most of its biomass (∼70%) remained at 16 h (Figure 3E, Table 1). The level of *vpsL-lux* expression in the Δ*rbmB* mutant was similar to the WT, showing that the RbmB dispersal function does not concern production of VPS (Figure 3F). Complementation with inducible *rbmB* expressed from an ectopic locus in the Δ*rbmB* strain caused a ∼40% reduction in peak biofilm formation, confirming that RbmB negatively regulates biofilm formation, however the complemented strain retained a modest biofilm dispersal defect, suggesting that the timing or level of *rbmB* expression is critical for WT biofilm disassembly (Supplementary Figure 3B). To verify that the Δ*rbmB* dispersal defect stems from the lack of *vps* degradation, we grew Δ*rbmB* biofilms for 16 h (i.e., post WT biofilmdispersal completion), and subsequently fixed and stained the non-dispersed biofilms with wheat germ agglutinin conjugated to Texas Red (WGA-txRed), which binds to N-acetylglucosamine sugars in the VPS matrix.(27) We used the Δ*lapG* mutant as our control since its biofilm dispersal phenotype should not involve changes in VPS. On average, the Δ*rbmB* mutant exhibited ∼6x more WGA-txRed signal than the Δ*lapG* mutant (Figure 3G). Collectively, our results show that the non-dispersed Δ*lapG* biofilms contain little VPS, consistent with possession of functional RbmB, while non-dispersed Δ*rbmB* biofilms contain excess VPS due to the lack of RbmB-mediated polysaccharide digestion. Thus, we suggest that RbmB-directed VPS disassembly is critical for proper biofilm disassembly (Figure 3H). Our results show that LapG and RbmB function in different pathways to drive biofilm disassembly. To examine their combined effects, we constructed the Δ*lapG* Δ*rbmB* double mutant and measured its biofilm lifecycle (Figure 3I). The Δ*lapG* Δ*rbmB* double mutant mimicked the single Δ*rbmB* mutant (Figure 3E) in its biofilm dispersal defect. Thus, the Δ*lapG* and Δ*rbmB* defects are not additive. Presumably, the severe dispersal defect displayed by the Δ*rbmB* single mutant, which cannot digest matrix polysaccharides, is not made more extreme by additional impairment of matrix protein degradation, suggesting that cells are already maximally trapped by the undigested polysaccharides.

Extracellular DNA (eDNA) is a component of the *V. cholerae* biofilm matrix and two DNAses secreted by *V. cholerae*, Dns and Xds, digest eDNA.(28) Although we did not identify *dns* and *xds* in our screen, we nonetheless investigated whether they contributed to biofilm dispersal. Neither the Δ*dns* and the Δ*xds* single mutants, nor the Δ*dns* Δ*xds* double mutant displayed a biofilm dispersal defect in our assay (Supplementary Figure 3C), suggesting that eDNA digestion is not required for dispersal. In a similar vein, we did not identify genes encoding the eight *V. cholerae* extracellular proteases that could degrade matrix proteins. Consistent with this finding, measurement of the phenotypes of mutants deleted for each extracellular protease gene showed that none exhibited a dispersal defect. Thus, no single extracellular protease is required for biofilm dispersal (Supplementary Figure 3D). It remains possible that proteases contribute to biofilm dispersal by functioning redundantly. Together, our results indicate that two enzymes, LapG and RbmB, are the primary matrix degrading components that enable biofilm dispersal.

### Reorientations in swimming direction are required for biofilm dispersal

The final category of genes identified in our screen are involved in cell motility. As noted above, non-motile mutants were excluded from analysis because they are known to be impaired in surface attachment. Nonetheless, we identified a mutant containing a transposon insertion in *cheY3* as defective for biofilm dispersal. *cheY3* is one of the five *V. cholerae cheY* genes specifying chemotaxis RR proteins.(29) Notably, *cheY3* is the only *V. cholerae cheY* homolog required for chemotaxis.(29) The Δ*cheY3* mutant exhibited similar peak biofilm timing and biomass as WT *V. cholerae*, however, ∼21% biomass remained at 16 h (Figure 4A, Table 1). Complementation via introduction of *cheY3* at an ectopic locus restored biofilm dispersal in the mutant (Supplementary Figure 4A). Expression of *vpsL-lux* in the Δ*cheY3* mutant was identical to the WT indicating that the dispersal phenotype was not due to elevated matrix production (Figure 4B).

**Figure 4.**
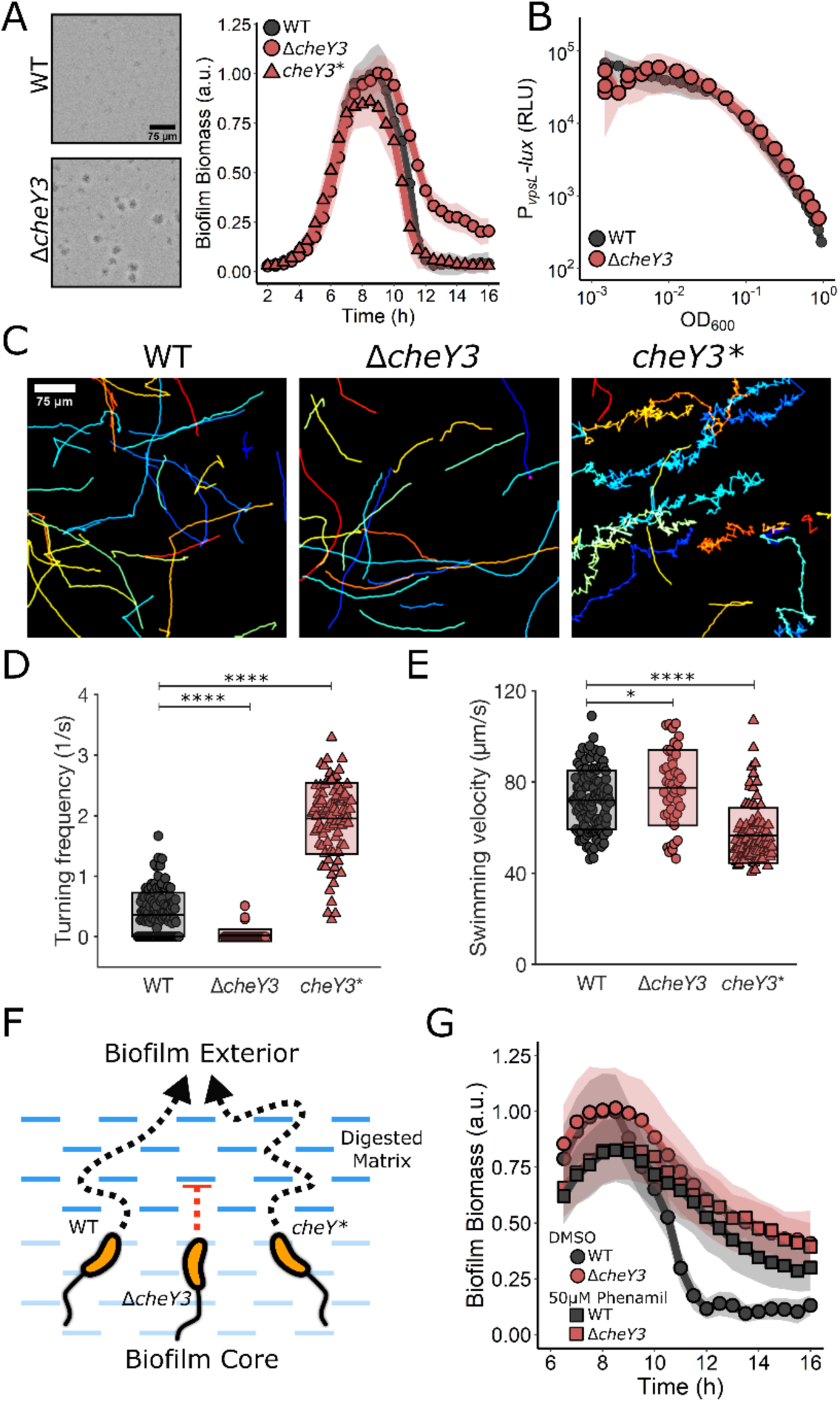
Reorientations in swimming direction are required for *V. cholerae* biofilm dispersal. (A) Representative 16 h images and quantitation of biofilm biomass over time measured by time-lapse microscopy for WT *V. cholerae*, the Δ*cheY3* mutant, and the *cheY3*^*D16K, Y109W*^ (*cheY3*\*) mutant. (B) The corresponding *P*_*vpsL*_*-lux* output for WT and the Δ*cheY3* strain over the growth curve. (C) Representative, randomly colored, single-cell locomotion trajectories for the strains in A. (D) Turning frequencies of the strains in A. (E) Measured swimming velocities of the strains in A. (F) Proposed model for the role of motility and reorientation in biofilm dispersal. (G) Quantitation of biofilm biomass over time for WT and the Δ*cheY3* mutant following treatment with DMSO or the motility inhibitor, phenamil supplied at 5 h post-inoculation. For biofilm biomass assays, *N* = 3 biological and *N* = 3 technical replicates, ± SD (shaded). a.u., arbitrary unit. For *vpsL-lux* measurements, *N* = 3 biological replicates, ± SD (shaded). RLU, relative light units. For motility measurements, 45-125 individual cells of each strain were tracked. In panels D and E, unpaired t-tests were performed for statistical analysis, with P values denoted as *P < 0.05; **P < 0.01; *** P < 0.001; ****P < 0.0001; n.s., P > 0.05.

The *V. cholerae* default motor rotation direction is counterclockwise (CCW), which fosters smooth, straight swimming.(30) Transition to clockwise (CW) motor rotation causes reorientations in swimming direction.(30) Phospho-CheY3 binds to the flagellar motor switch complex to mediate the change from CCW to CW rotation. Thus, the *ΔcheY3* mutant is non-chemotactic and the cells are locked in the CCW, straight swimming mode (Figure 4C). We reasoned that the *ΔcheY3* mutant dispersal defect could stem from an inability to chemotact or from an inability to reorient swimming direction. To distinguish between these possibilities, we examined biofilm dispersal in a *V. cholerae* mutant carrying a *cheY3* allele, *cheY3*^*D16K, Y109W*^ (henceforth, *cheY3\**) that locks the motor into CW rotation and so also disrupts chemotaxis. *cheY3\** cells undergo frequent reorientations and are unable to swim in smooth straight runs (Figure 4C).(29, 31) The *cheY3\** strain had WT biofilm dispersal capability. Thus, being chemotactic is not required for *V. cholerae* to exit biofilms (Figure 4A).

We reasoned that analysis of the unique motility characteristics of our strains could reveal the underlying causes of the *ΔcheY3* biofilm dispersal defect. We measured the turning frequencies and swimming velocities of the WT, *ΔcheY3*, and *cheY3* V. cholerae* strains. Consistent with previous reports, these three mutants exhibited notable differences: on average, the WT turned once every 3 s, the *ΔcheY3* mutant turned less than once every 40 s, and the *cheY3\** strain turned once every 0.5 s (Figure 4C and D).(29, 31) The *cheY3\** strain displayed slightly lower average swimming velocity than the WT and *ΔcheY3* strains, due to its high turning frequency as turning necessarily involves a decrease in velocity (Figure 4E).(32) Together, these results suggest that the low turning frequency of the *ΔcheY3* mutant is responsible for the biofilm dispersal defect. We propose that if cells do not frequently change their direction of motion, they become trapped by the biofilm matrix mesh which compromises their ability to escape (Figure 4F). Indeed, in other bacteria, straight-swimming mutants are deficient in traversing fluid-filled porous media compared to WT organisms that can reorient.(33) Together, these results indicate that chemotaxis itself is not required for biofilm dispersal, but, rather, that the chemotaxis machinery facilitates random reorientation events that allow *V. cholerae* cells to navigate a porous biofilm matrix. The same non-chemotactic mutants used here exhibit stark differences in competition experiments in animal models of cholera infection, showing that their differences in motility and, possibly, their differences in biofilm dispersal capabilities, are pertinent to colonization.(31)

Finally, we determined whether the ability to locomote was required for biofilm dispersal or, by contrast, if non-motile cells could escape the digested matrix via Brownian motion. As mentioned above, we could not simply study dispersal of non-flagellated and non-motile mutants because of their confounding surface attachment defects and feedback on biofilm regulatory components.(34, 35) To circumvent this problem, we employed phenamil, an inhibitor of the Na^+^-driven *V. cholerae* flagellar motor, which, as expected, dramatically reduced planktonic cell motility (Supplementary Figure 4B).(36) To assess the role of swimming motility in biofilm dispersal, we first allowed WT *V. cholerae* cells to undergo biofilm formation for 5 h, at which point we perfused DMSO or phenamil into the incubation chamber (Figure 4G). Following phenamil treatment, the WT strain displayed a dispersal defect nearly identical to that of the *ΔcheY3* mutant. Additionally, phenamil treatment of the *ΔcheY3* mutant did not further impair its biofilm dispersal. Together, these results demonstrate that swimming motility is crucial for *V. cholerae* biofilm dispersal and an inability to reorient is as detrimental to dispersal as a complete lack of flagellar motility.

## Discussion

In this study, we developed a high-content imaging screen that allowed us to identify components required for *V. cholerae* biofilm dispersal. We categorized the identified components into three classes: signal transduction, matrix disassembly, and cell motility. We propose that the three functional categories represent the chronological steps required for the disassembly of a biofilm: First, the stimuli that activate dispersal must accumulate. Subsequently, the gene expression pattern established by detection of these stimuli must repress biofilm matrix production and activate production of enzymes required to digest the biofilm matrix. Finally, cells must escape through the partially digested, porous matrix which requires changes in the direction of movement. Together, these steps ensure that when environmental conditions are appropriate, *V. cholerae* cells can exit the sessile lifestyle and disseminate to new terrain that is ripe for biofilm formation or, alternatively, during disease, to a new host. One can now imagine targeting the functions identified in this work for small-molecule disruption of the *V. cholerae* biofilm lifecycle, possibly guiding the development of treatments to reduce the duration of *V. cholerae* infection or to prevent transmission.

## Materials and Methods

### Bacterial Strains and Reagents

The *V. cholerae* parent strain used in this study was WT O1 El Tor biotype C6706str2. Antibiotics were used at the following concentrations: polymyxin B, 50 μg/mL; kanamycin, 50 μg/mL; spectinomycin, 200 μg/mL; and chloramphenicol, 1 μg/mL. Strains were propagated in lysogeny broth (LB) supplemented with 1.5% agar or in liquid LB with shaking at 30°C. All strains used in this work are reported in Supplementary Table 1. Unless otherwise stated, exogenous compounds were added from the onset of biofilm initiation. The antimicrobial peptide C18G (VWR) was added at 5 µg/mL. Phenamil (Sigma) was prepared in DMSO and added 5 h post biofilm inoculation to a final concentration of 50 µM. L-arabinose (Sigma) was prepared in water and added at 0.2%.

### DNA Manipulation and Strain Construction

To produce linear DNA fragments for natural transformations, splicing overlap extension PCR was performed using iProof polymerase (Bio-Rad, Hercules, CA, USA) to combine DNA pieces. Primers and gene fragments used in this study are reported in Supplementary Table 2. In all cases, ∼3 kb of upstream and downstream flanking regions of homology were generated by PCR from *V. cholerae* genomic DNA and were included to ensure high chromosomal integration frequency. DNA fragments that were not native to *V. cholerae* were synthesized as g-blocks (IDT, Coralville, IA, USA).

All *V. cholerae* strains generated in this work were constructed by replacing genomic DNA with DNA introduced by natural transformation as previously described.(18, 37) The neutral *vc1807* locus was used as the site of introduction of the gene encoding the antibiotic resistance cassette in the natural co-transformation procedure. The *vc1807* locus was also used as the site for introduction of genes under study in chromosomal ectopic expression analyses.(37) PCR and Sanger sequencing were used to verify correct integration events. Genomic DNA from recombinant strains was used for future co-transformations and as templates for PCR to generate DNA fragments, when necessary. Deletions were constructed in frame and eliminated the entire coding sequences. The exceptions were *mbaA, dbfS*, and *dbfR*, which each overlap with another gene in their operons. In these cases, portions of the genes were deleted ensuring that adjacent genes were not perturbed. For *tagA*, the first 103 base pairs, including the nucleotides specifying the start codon, were deleted. All strains constructed in this study were verified by sequencing at Genewiz.

### Microscopy and Mutant Screening

The biofilm lifecycle was measured using time-lapse microscopy as described previously.(18) All plots were generated using ggplot2 in R. To generate the library of *V. cholerae* insertion mutants for the dispersal screen, the WT parent strain was mutagenized with Tn*5* as previously described.(38) Mutants were selected by growth overnight on LB plates containing polymyxin B and kanamycin. The next day, mutant colonies were arrayed into 96-well plates containing 200 µL of LB medium supplemented with polymyxin B and kanamycin using an automated colony-picking robot (Molecular Devices). The arrayed cultures were grown in a plate-shaking incubator at 30^°^ C covered with breathe-easy membranes to minimize evaporation. After 16 h of growth, the arrayed cultures were diluted 1:200,000 into 96-well plates containing M9 medium supplemented with glucose and casamino acids. Diluted cultures were incubated statically at 30^°^ C for 8 h (to achieve peak biofilm biomass), at which point, images of each well were captured on a Nikon Ti-E inverted microscope using transmitted-light bright-field illumination, a 10× Plan Fluor (NA 0.3) objective lens, and an Andor iXon 897 EMCCD camera. Automated image acquisition was performed using NIS-Elements software v5.11.02 and the NIS-Elements Jobs Module to acquire images at four positions within each well to account for heterogeneity within samples. To maintain the focal plane between wells, the Nikon Perfect Focus System was used. After performing microscopy at the 8 h timepoint, 96-well plates were returned to the incubator. To assess biofilm dispersal, a second set of images of the same samples was acquired at 13 h post inoculation. Mutants that displayed biofilm growth at the 8 h timepoint but failed to disperse by the 13 h timepoint were subcultured, grown overnight, and subsequently re-imaged using the time-lapse approach described above to assess their biofilm lifecycles in real-time. Mutants that exhibited biofilm dispersal defects after this reassessment step were analyzed for the locations of transposon insertions using arbitrary PCR.(39)

### *lux* Transcription Assays

Three colonies of each strain to be analyzed were individually grown overnight in 200 μL LB with shaking at 30°C in a 96-well plate covered with a breathe-easy membrane. The following morning, the cultures were diluted 1:5,000 into fresh M9 medium supplemented with glucose and casamino acids. The plates were placed in a BioTek Synergy Neo2 Multi-Mode reader (BioTek, Winooski, VT, USA) under static growth conditions at 30°C. Both OD_600_ and bioluminescence from the *lux* fusions were simultaneously measured at 15 min time intervals. Results were exported to R, and light values were divided by OD_600_ to produce relative light units (RLUs). Results from replicates were averaged and plotted using ggplot2 in R.

### VPS Quantitation

To assess VPS levels in non-dispersed biofilms using WGA-txRED, biofilms were grown for 16 h and subsequently washed 3 times with 1× phosphate buffered saline (PBS), and fixed for 10 min with 3.7% formaldehyde in 1× PBS. After fixation, samples were washed 5 times with 1× PBS and subsequently incubated with a solution containing 1 µg/mL WGA-txRED (ThermoFisher Scientific), 1 µg/mL 4’, 6-diamidino-2-phenylindole (DAPI), and 1% bovine serum albumin in 1× PBS for 1 h with shaking at 30^°^ C in the dark. After incubation, samples were washed 5 more times with 1× PBS before imaging. Confocal microscopy was performed on a Leica DMI8 SP-8 point scanning confocal microscope (Leica, Wetzlar, Germany) with the pinhole set to 1.0 airy unit. The light source for DAPI was a 405 laser and the light source used to excite WGA-txRED was a tunable white-light laser (Leica; model #WLL2; excitation window = 470–670 nm) set to 595 nm. Biofilms were imaged using a 10× air objective (Leica, HC PL FLUOTAR; NA: 0.30). Sequential frame scanning was performed to minimize spectral bleed-through in images. Emitted light was detected using GaAsP spectral detectors (Leica, HyD SP), and timed gate detection was employed to minimize the background signal. Image analyses were performed in FIJI software (Version 1.52p). Biofilms were segmented in the DAPI channel using an intensity threshold and the intensities of each channel were measured. The same threshold was applied to all images. WGA-txRED signal was divided by DAPI signal to achieve the normalized WGA signal.

### Motility Assay

To prevent biofilm formation during measurements of swimming velocities and turning frequencies for the WT, Δ*cheY3*, and *cheY3\** strains, *vpsL* was deleted. Each strain was grown for 16 h in LB medium and the following day, cells were diluted to OD_600_ = 0.001 in M9 medium supplemented with glucose and casamino acids. Subsequently, diluted cultures were dispensed in 200 µL aliquots into glass-coverslip bottomed 96-well plates (MatTek, Ashland, MA, USA). After a period of 1 h, during which time cells were allowed to adhere to the coverslips, wells were washed 8 times with fresh medium to remove unattached cells. The plates were incubated at 25^°^ C for 3 h, and imaging was performed using the brightfield setup described above for the biofilm dispersal screen. In this case, the frame interval was 50 msec and imaging was conducted at a distance of ∼100 µm into the sample. Images were smoothed, background corrected, and imported into the TrackMate (v.5.2.0) plugin in FIJI. Cells were detected with a Laplacian of Gaussian (LoG) detector and were subsequently tracked using the simple Linear Assignment Problem (LAP) approach. To exclude non-motile cells from our analyses in Figure 4C-E objects with velocities under 40 µm/sec were eliminated. Analyses and plotting of swimming velocities and turning frequencies were performed in MATLAB (The Mathworks, Inc.). Local curvatures for single-cell locomotion trajectories were calculated as described.(40) Curvature of less than 0.3 μm^-1^ was used to identify the turning events. MSD was calculated as described previously.(41)

### Phos-tag Gel Analysis

To monitor DbfR and phospho-DbfR via SDS-PAGE, the endogenous *dfbR* gene was replaced with *dbfR-SNAP* in the *ΔdbfS* strain, and *P*_*BAD*_*-dbfS* was introduced at the ectopic locus, *vc1807*. To assess DbfR-SNAP phosphorylation in the absence and presence of DbfS, overnight cultures of the strain were diluted 1:1000 and subsequently grown for 4 h at 30^°^ C with shaking to an OD_600_ ∼ 0.6. To each culture, 1 µM SNAP-Cell TMR Star (New England Biolabs) was added to label the SNAP tag, and the culture was subsequently divided into two tubes. To one tube, 0.2% D-fucose was added, and to the other, 0.2% L-arabinose was added to repress and induce DbfS production, respectively. The cultures were returned to 30^°^ C with shaking. After 1 h, the cells were collected by centrifugation for 1 min at 13,000 rpm. Lysis and solubilization were carried out as rapidly as possible. Briefly, cells were chemically lysed by resuspension to OD_600_ = 1.0 in 40 μL Bug Buster (Novagen) for 5 min at 25° C with intermittent vortex. The cell lysate was solubilized at 25^°^ C in 1.5× SDS-PAGE buffer for 5 min also with intermittent vortex. Samples were immediately loaded onto a cold 7.5% SuperSep™ Phos-tag™ (50 μM/L) gel (FUJIFILM Wako Pure Chemical, 198-17981). Electrophoresis was carried out at 100 V at 4^°^ C until the loading buffer exited the gel. Gel images were captured on an ImageQuant LAS 4000 imager (GE Healthcare) using a Cy3 filter set.

---

**Supplementary Figure 1.**
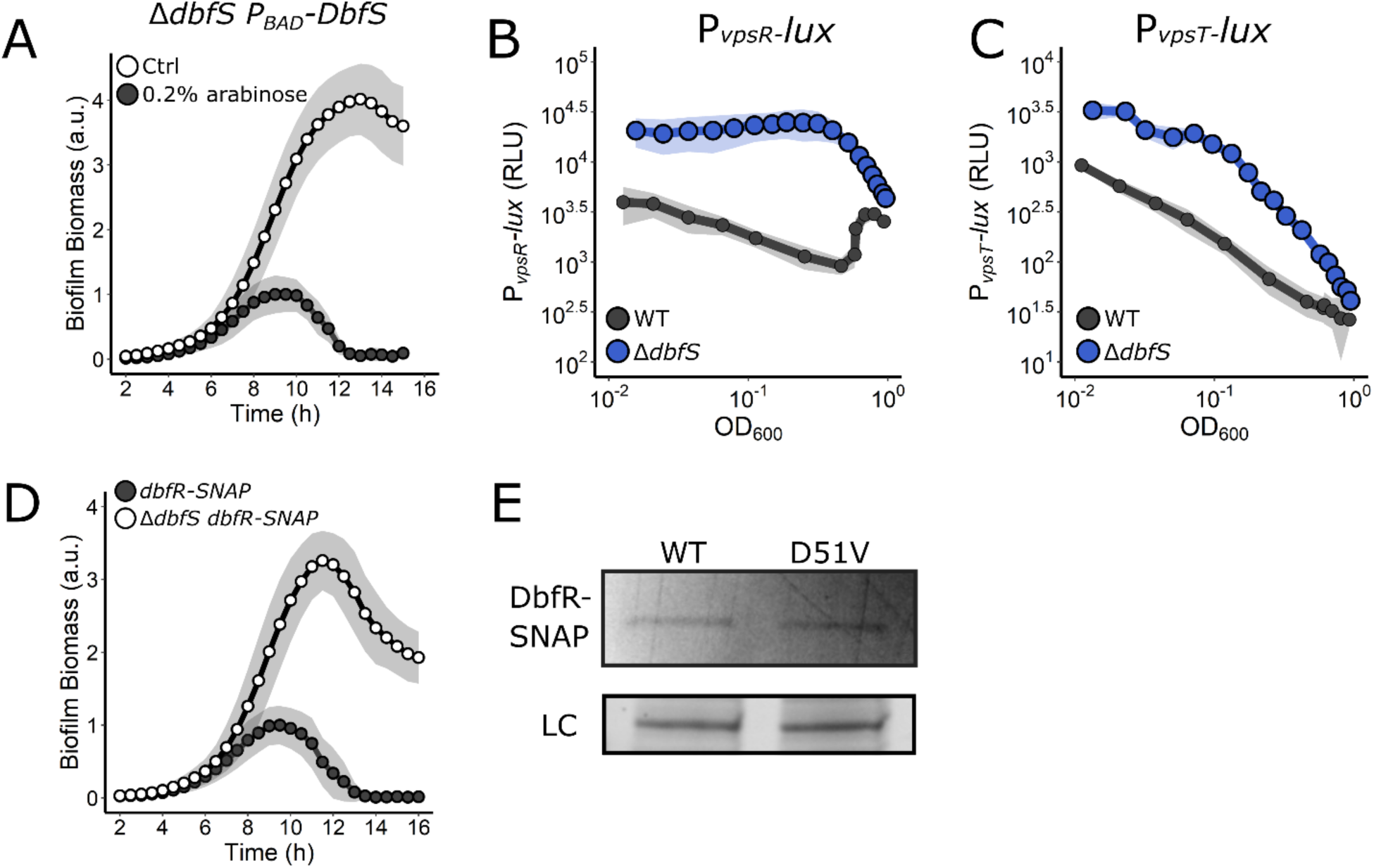
Complementation, functional tagging, and mutagenesis of the DbfS-DbfR two-component system. (A) Quantitation of biofilm biomass over time measured by time-lapse microscopy for the Δ*dbfS P*_*BAD*_*-dbfS* strain following addition of water (Ctrl) or 0.2% arabinose. (B) *P*_*vpsR*_*-lux* and (C) *P*_*vpsT*_*-lux* output for WT and the Δ*dbfS* strain over the growth curve. (D) As in A for SNAP-tagged DbfR in the WT and Δ*dbfS* strains. (E) Top panel: representative in-gel SDS-PAGE fluorescence following electrophoresis of *V. cholerae* cell lysates containing WT DbfS-SNAP or DbfS^D51V^-SNAP that had been incubated with SNAP-Cell TMR Star. Bottom panel: Coomassie stained loading control (LC). For all biofilm measurements, *N* = 3 biological and *N* = 3 technical replicates, ± SD (shaded). a.u., arbitrary unit. For *lux* measurements, *N* = 3 biological replicates, ± SD (shaded). RLU, relative light units.

**Supplementary Figure 2.**
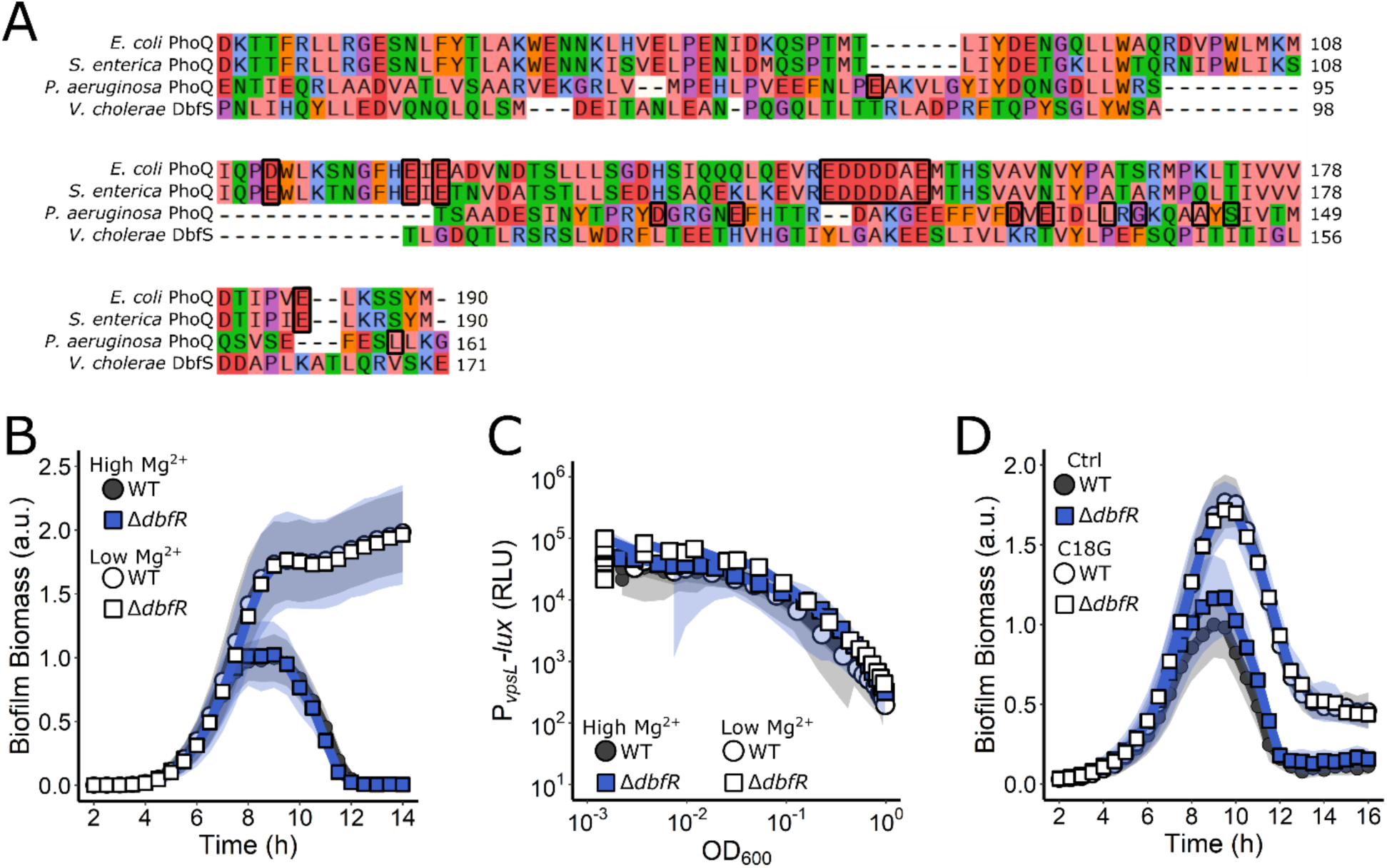
DbfS is not functionally equivalent to PhoQ. (A) Alignment of the sensory domains of PhoQ from *E. coli, S. enterica*, and *P. aeruginosa* against that of *V. cholerae* DbfS. Black boxes indicate residues involved in Mg^2+^ binding in PhoQ. (B) Quantitation of biofilm biomass over time measured by time-lapse microscopy in high magnesium (10 mM) and limiting magnesium (10 µM) conditions for WT *V. cholerae* and the Δ*dbfR* strain. (C) The corresponding *P*_*vpsL*_*-lux* outputs for strains and growth conditions in B over the growth curve. (D) As in B except following the addition of water or 5 µg/mL C18G. In all cases, *N* = 3 biological and *N* = 3 technical replicates, ± SD (shaded). a.u., arbitrary unit. For *vpsL-lux* measurements, *N* = 3 biological replicates, ± SD (shaded). RLU, relative light units.

**Supplementary Figure 3.**
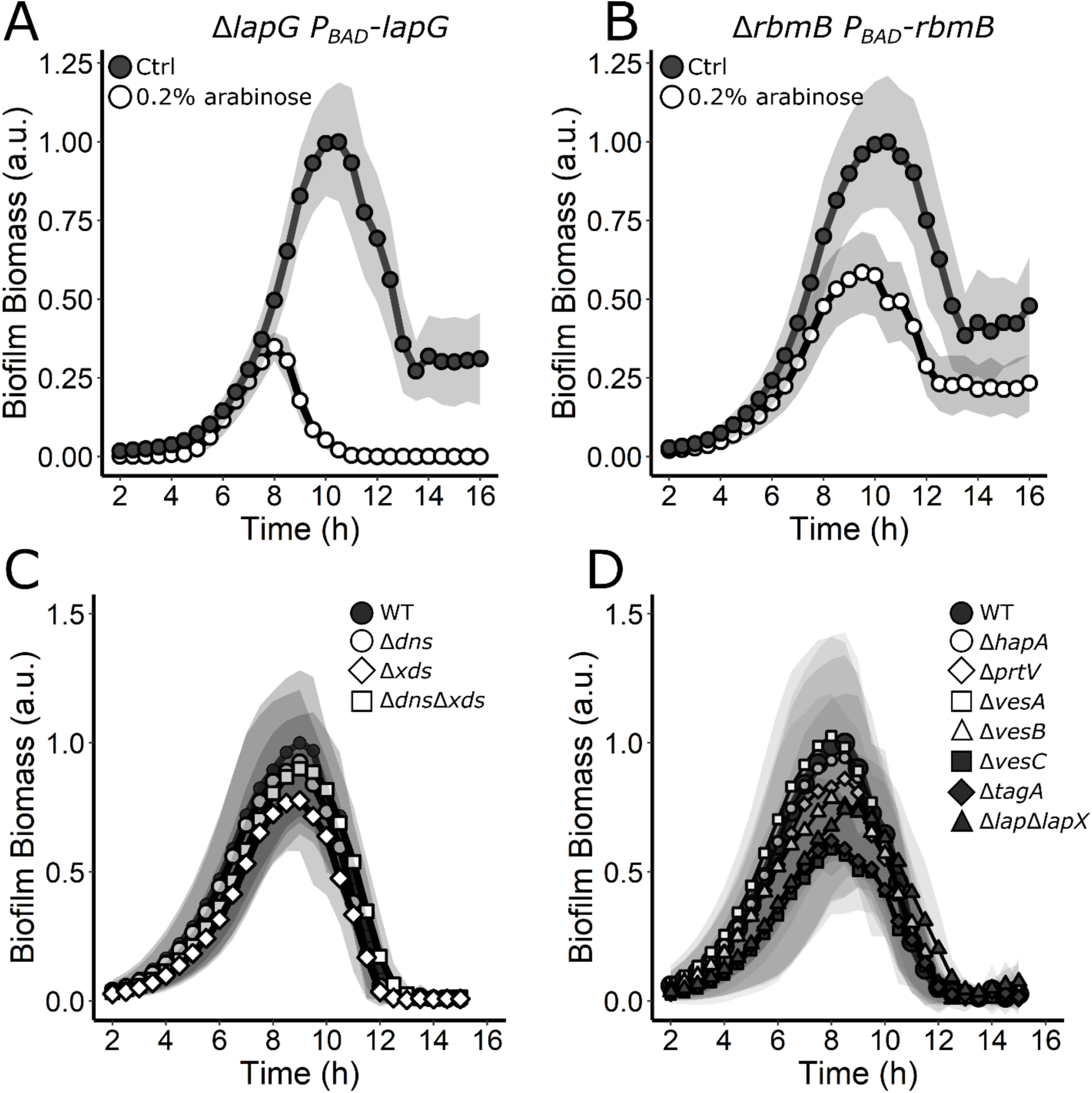
Introduction of *lapG* and *rbmB* complements the Δ*lapG* and Δ*rbmB* biofilm defects, respectively, and assessment of the roles of extracellular DNAses and secreted proteases in *V. cholerae* biofilm dispersal. (A) Quantitation of biofilm biomass over time measured by time-lapse microscopy for the Δ*lapG P*_*BAD*_*-lapG* strain following addition of water (Ctrl) or 0.2% arabinose. (B) As in A, but for the Δ*rbmB P*_*BAD*_*-rbmB* strain. (C) Quantitation of biofilm biomass over time measured by time-lapse microscopy for WT *V. cholerae* and mutants lacking the designated DNAses. (D) Quantitation of biofilm biomass over time measured by time-lapse microscopy for WT *V. cholerae* and mutants lacking the designated proteases. In all cases, *N* = 3 biological and *N* = 3 technical replicates, ± SD (shaded). a.u., arbitrary unit.

**Supplementary Figure 4.**
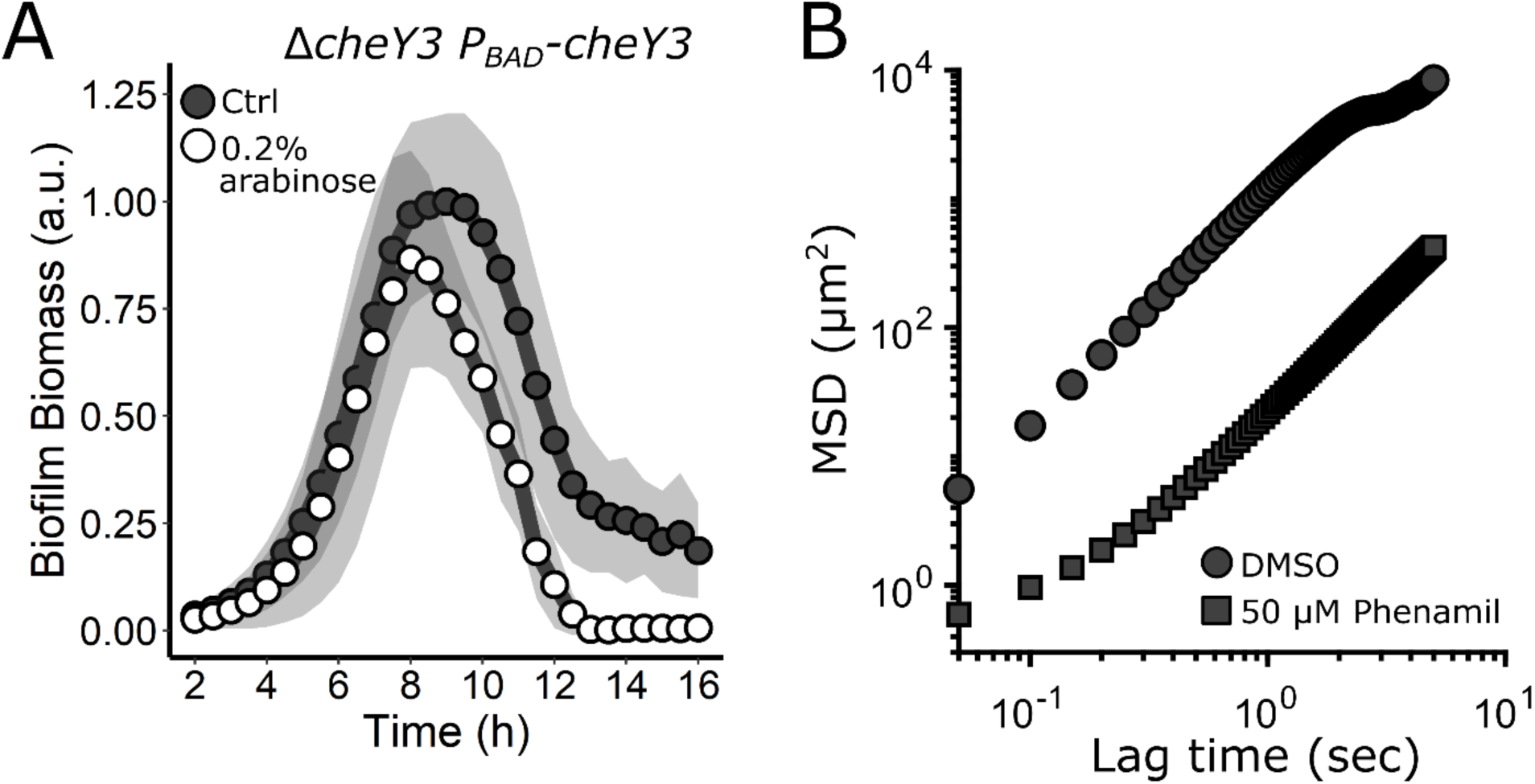
Complementation of the Δ*cheY3* mutant and phenamil inhibition of *V. cholerae* motility. (A) Quantitation of biofilm biomass over time measured by time-lapse microscopy for the Δ*cheY3 P*_*BAD*_*-cheY3* strain following addition of water (Ctrl) or 0.2% arabinose. In all cases, *N* = 3 biological and *N* = 3 technical replicates, ± SD (shaded). a.u., arbitrary unit. (B) Mean squared displacement (MSD) of cell trajectories versus lag time for WT *V. cholerae* treated with DMSO solvent or 50 µM phenamil.

## Supplemental Discussion

### DbfS is not equivalent to PhoQ

In *E. coli*, low Mg^2+^ and cationic peptides activate PhoQ kinase activity.(42) Sequence alignment of the DbfS sensory domain with that from PhoQ of *E. coli, Salmonella enterica*, and *Pseudomonas aeruginosa* revealed that DbfS lacks all of the key residues involved in Mg^2+^ binding (Supplementary Figure 2A).(43) To test if Mg^2+^ alters DfbS activity, we measured the *V. cholerae* biofilm lifecycle in response to low Mg^2+^ conditions in WT *V. cholerae* and in the Δ*dbfR* mutant. If, analogous to PhoQ, DfbS kinase activity is activated by low Mg^2+^, when Mg^2+^ is limiting, WT *V. cholerae* should exhibit an altered biofilm dispersal phenotype while the Δ*dbfR* mutant would be impervious to Mg^2+^ changes.(42) Supplementary Figure 2B shows that Mg^2+^ limitation does indeed inhibit *V. cholerae* biofilm dispersal, however, inhibition occurs in *both* the WT and the Δ*dbfR* strains. Mg^2+^ limitation did not alter *vpsL-lux* expression in either strain (Supplementary Figure 2C). Thus, Mg^2+^ does not control DfbS activity. We obtained the same results following exogenous addition of the cationic peptide C18G (Supplementary Figure 2D). Together, these results demonstrate that DfbS does not respond to the ligands that control PhoQ activity.

## Acknowledgements

We thank members of the Bassler group and Prof. Ned Wingreen for thoughtful discussions. We particularly thank Dr. Matthew Jemielita for help with the secreted protease mutants used in this study. This work was supported by the Howard Hughes Medical Institute, NIH Grant 5R37GM065859, National Science Foundation Grant MCB-1713731, and a Max Planck-Alexander von Humboldt research award to BLB. AAB is a Howard Hughes Medical Institute Fellow of the Damon Runyon Cancer Research Foundation, DRG-2302-17. The content is solely the responsibility of the authors and does not necessarily represent the official views of the National Institutes of Health. The funders had no role in study design, data collection and analysis, decision to publish, or preparation of the manuscript.

The authors declare no competing interest.

---

**Supplementary Table 1.**
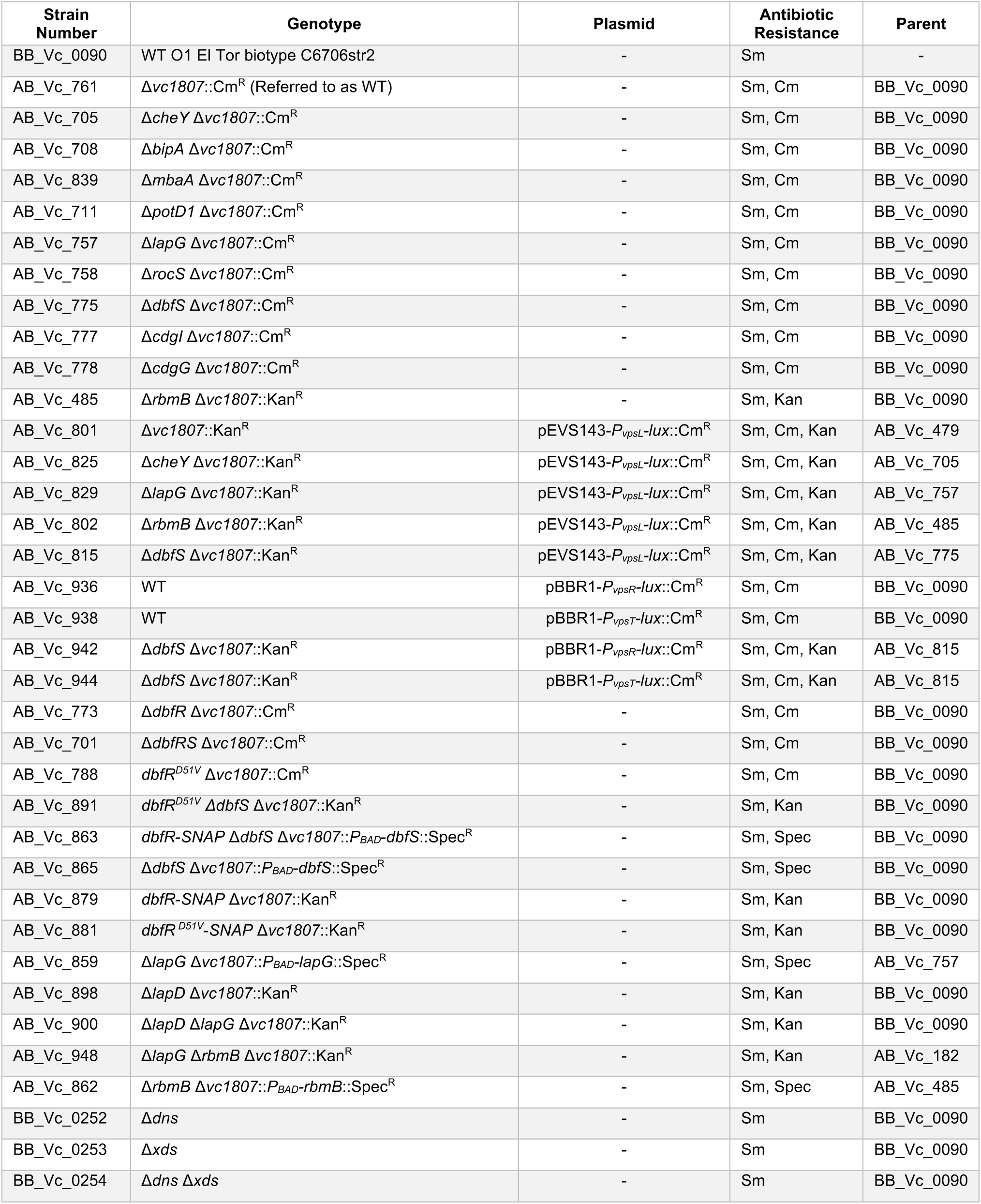
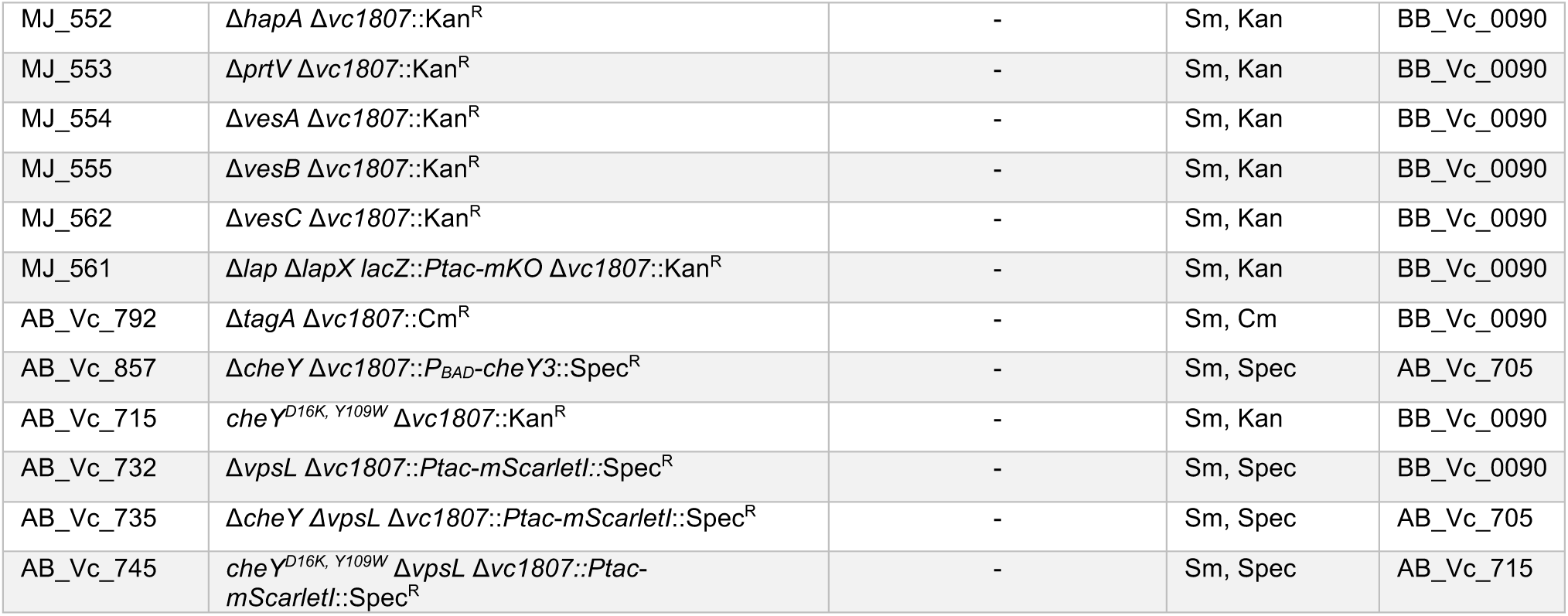
Strains used in this study.

| Strain Number | Genotype | Plasmid | Antibiotic Resistance | Parent |
| --- | --- | --- | --- | --- |
| BB_Vc_0090 | WT O1 El Tor biotype C6706str2 | - | Sm | - |
| AB_Vc_761 | $\Delta vc1807::Cm^R$ (Referred to as WT) | - | Sm, Cm | BB_Vc_0090 |
| AB_Vc_705 | $\Delta cheY \Delta vc1807::Cm^R$ | - | Sm, Cm | BB_Vc_0090 |
| AB_Vc_708 | $\Delta bipA \Delta vc1807::Cm^R$ | - | Sm, Cm | BB_Vc_0090 |
| AB_Vc_839 | $\Delta mbaA \Delta vc1807::Cm^R$ | - | Sm, Cm | BB_Vc_0090 |
| AB_Vc_711 | $\Delta potD1 \Delta vc1807::Cm^R$ | - | Sm, Cm | BB_Vc_0090 |
| AB_Vc_757 | $\Delta lapG \Delta vc1807::Cm^R$ | - | Sm, Cm | BB_Vc_0090 |
| AB_Vc_758 | $\Delta rocS \Delta vc1807::Cm^R$ | - | Sm, Cm | BB_Vc_0090 |
| AB_Vc_775 | $\Delta dbfS \Delta vc1807::Cm^R$ | - | Sm, Cm | BB_Vc_0090 |
| AB_Vc_777 | $\Delta cdgI \Delta vc1807::Cm^R$ | - | Sm, Cm | BB_Vc_0090 |
| AB_Vc_778 | $\Delta cdgG \Delta vc1807::Cm^R$ | - | Sm, Cm | BB_Vc_0090 |
| AB_Vc_485 | $\Delta rbmB \Delta vc1807::Kan^R$ | - | Sm, Kan | BB_Vc_0090 |
| AB_Vc_801 | $\Delta vc1807::Kan^R$ | pEVS143-$P_{vpsL}$-$lux::Cm^R$ | Sm, Cm, Kan | AB_Vc_479 |
| AB_Vc_825 | $\Delta cheY \Delta vc1807::Kan^R$ | pEVS143-$P_{vpsL}$-$lux::Cm^R$ | Sm, Cm, Kan | AB_Vc_705 |
| AB_Vc_829 | $\Delta lapG \Delta vc1807::Kan^R$ | pEVS143-$P_{vpsL}$-$lux::Cm^R$ | Sm, Cm, Kan | AB_Vc_757 |
| AB_Vc_802 | $\Delta rbmB \Delta vc1807::Kan^R$ | pEVS143-$P_{vpsL}$-$lux::Cm^R$ | Sm, Cm, Kan | AB_Vc_485 |
| AB_Vc_815 | $\Delta dbfS \Delta vc1807::Kan^R$ | pEVS143-$P_{vpsL}$-$lux::Cm^R$ | Sm, Cm, Kan | AB_Vc_775 |
| AB_Vc_936 | WT | pBBR1-$P_{vpsR}$-$lux::Cm^R$ | Sm, Cm | BB_Vc_0090 |
| AB_Vc_938 | WT | pBBR1-$P_{vpsT}$-$lux::Cm^R$ | Sm, Cm | BB_Vc_0090 |
| AB_Vc_942 | $\Delta dbfS \Delta vc1807::Kan^R$ | pBBR1-$P_{vpsR}$-$lux::Cm^R$ | Sm, Cm, Kan | AB_Vc_815 |
| AB_Vc_944 | $\Delta dbfS \Delta vc1807::Kan^R$ | pBBR1-$P_{vpsT}$-$lux::Cm^R$ | Sm, Cm, Kan | AB_Vc_815 |
| AB_Vc_773 | $\Delta dbfR \Delta vc1807::Cm^R$ | - | Sm, Cm | BB_Vc_0090 |
| AB_Vc_701 | $\Delta dbfRS \Delta vc1807::Cm^R$ | - | Sm, Cm | BB_Vc_0090 |
| AB_Vc_788 | $dbfR^{D51V} \Delta vc1807::Cm^R$ | - | Sm, Cm | BB_Vc_0090 |
| AB_Vc_891 | $dbfR^{D51V} \Delta dbfS \Delta vc1807::Kan^R$ | - | Sm, Kan | BB_Vc_0090 |
| AB_Vc_863 | $dbfR$-SNAP $\Delta dbfS \Delta vc1807::P_{BAD}$-$dbfS::Spec^R$ | - | Sm, Spec | BB_Vc_0090 |
| AB_Vc_865 | $\Delta dbfS \Delta vc1807::P_{BAD}$-$dbfS::Spec^R$ | - | Sm, Spec | BB_Vc_0090 |
| AB_Vc_879 | $dbfR$-SNAP $\Delta vc1807::Kan^R$ | - | Sm, Kan | BB_Vc_0090 |
| AB_Vc_881 | $dbfR^{D51V}$-SNAP $\Delta vc1807::Kan^R$ | - | Sm, Kan | BB_Vc_0090 |
| AB_Vc_859 | $\Delta lapG \Delta vc1807::P_{BAD}$-$lapG::Spec^R$ | - | Sm, Spec | AB_Vc_757 |
| AB_Vc_898 | $\Delta lapD \Delta vc1807::Kan^R$ | - | Sm, Kan | BB_Vc_0090 |
| AB_Vc_900 | $\Delta lapD \Delta lapG \Delta vc1807::Kan^R$ | - | Sm, Kan | BB_Vc_0090 |
| AB_Vc_948 | $\Delta lapG \Delta rbmB \Delta vc1807::Kan^R$ | - | Sm, Kan | AB_Vc_182 |
| AB_Vc_862 | $\Delta rbmB \Delta vc1807::P_{BAD}$-$rbmB::Spec^R$ | - | Sm, Spec | AB_Vc_485 |
| BB_Vc_0252 | $\Delta dns$ | - | Sm | BB_Vc_0090 |
| BB_Vc_0253 | $\Delta xds$ | - | Sm | BB_Vc_0090 |
| BB_Vc_0254 | $\Delta dns \Delta xds$ | - | Sm | BB_Vc_0090 |
| MJ_552 | *ΔhapA Δvc1807::Kan<sup>R</sup>* | - | Sm, Kan | BB_Vc_0090 |
| MJ_553 | *ΔprtV Δvc1807::Kan<sup>R</sup>* | - | Sm, Kan | BB_Vc_0090 |
| MJ_554 | *ΔvesA Δvc1807::Kan<sup>R</sup>* | - | Sm, Kan | BB_Vc_0090 |
| MJ_555 | *ΔvesB Δvc1807::Kan<sup>R</sup>* | - | Sm, Kan | BB_Vc_0090 |
| MJ_562 | *ΔvesC Δvc1807::Kan<sup>R</sup>* | - | Sm, Kan | BB_Vc_0090 |
| MJ_561 | *Δlap ΔlapX lacZ::Ptac-mKO Δvc1807::Kan<sup>R</sup>* | - | Sm, Kan | BB_Vc_0090 |
| AB_Vc_792 | *ΔtagA Δvc1807::Cm<sup>R</sup>* | - | Sm, Cm | BB_Vc_0090 |
| AB_Vc_857 | *ΔcheY Δvc1807::P<sub>BAD</sub>-cheY3::Spec<sup>R</sup>* | - | Sm, Spec | AB_Vc_705 |
| AB_Vc_715 | *cheY<sup>D16K, Y109W</sup> Δvc1807::Kan<sup>R</sup>* | - | Sm, Kan | BB_Vc_0090 |
| AB_Vc_732 | *ΔvpsL Δvc1807::Ptac-mScarletI::Spec<sup>R</sup>* | - | Sm, Spec | BB_Vc_0090 |
| AB_Vc_735 | *ΔcheY ΔvpsL Δvc1807::Ptac-mScarletI::Spec<sup>R</sup>* | - | Sm, Spec | AB_Vc_705 |
| AB_Vc_745 | *cheY<sup>D16K, Y109W</sup> ΔvpsL Δvc1807::Ptac-mScarletI::Spec<sup>R</sup>* | - | Sm, Spec | AB_Vc_715 |

**Supplementary Table 2.**
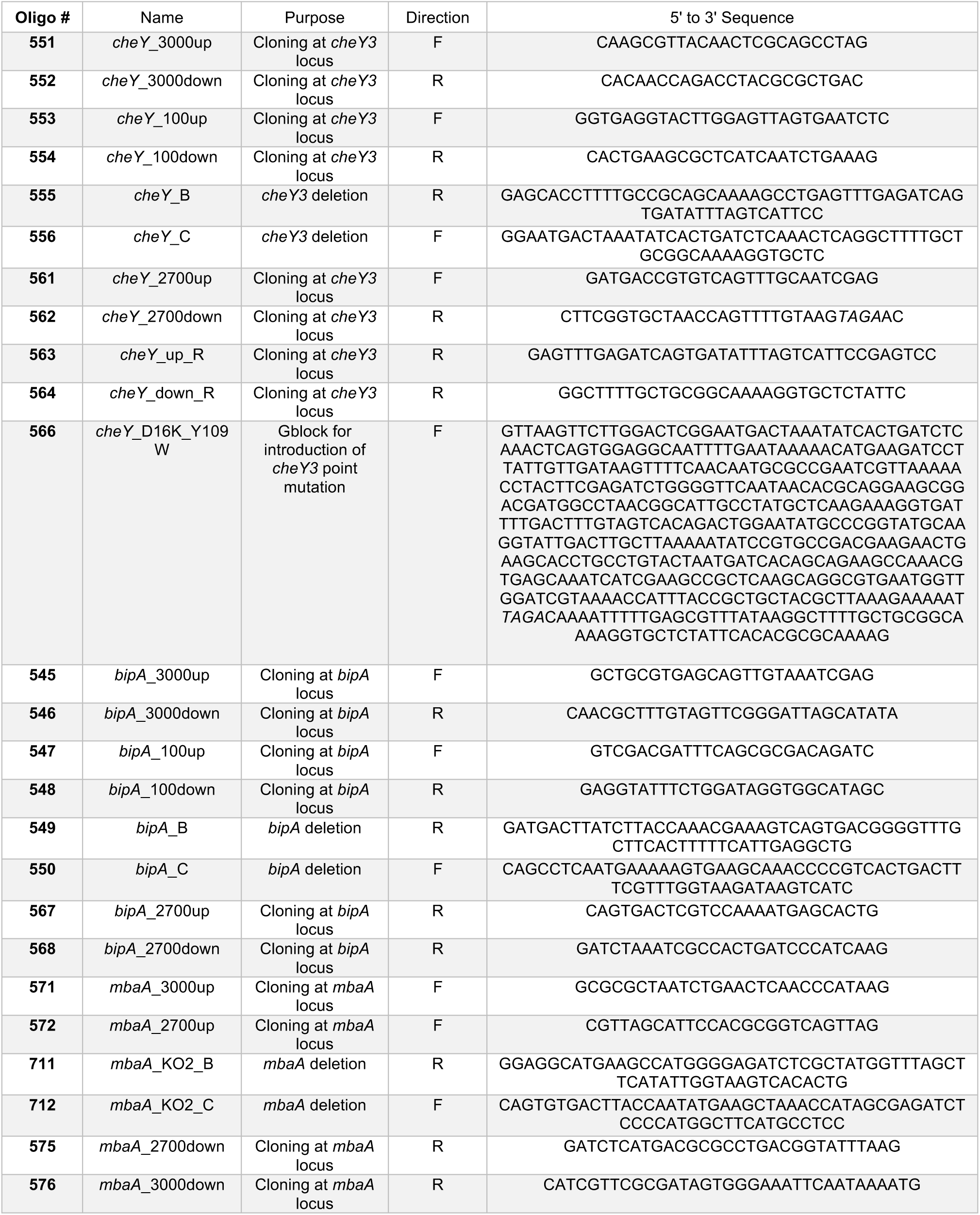
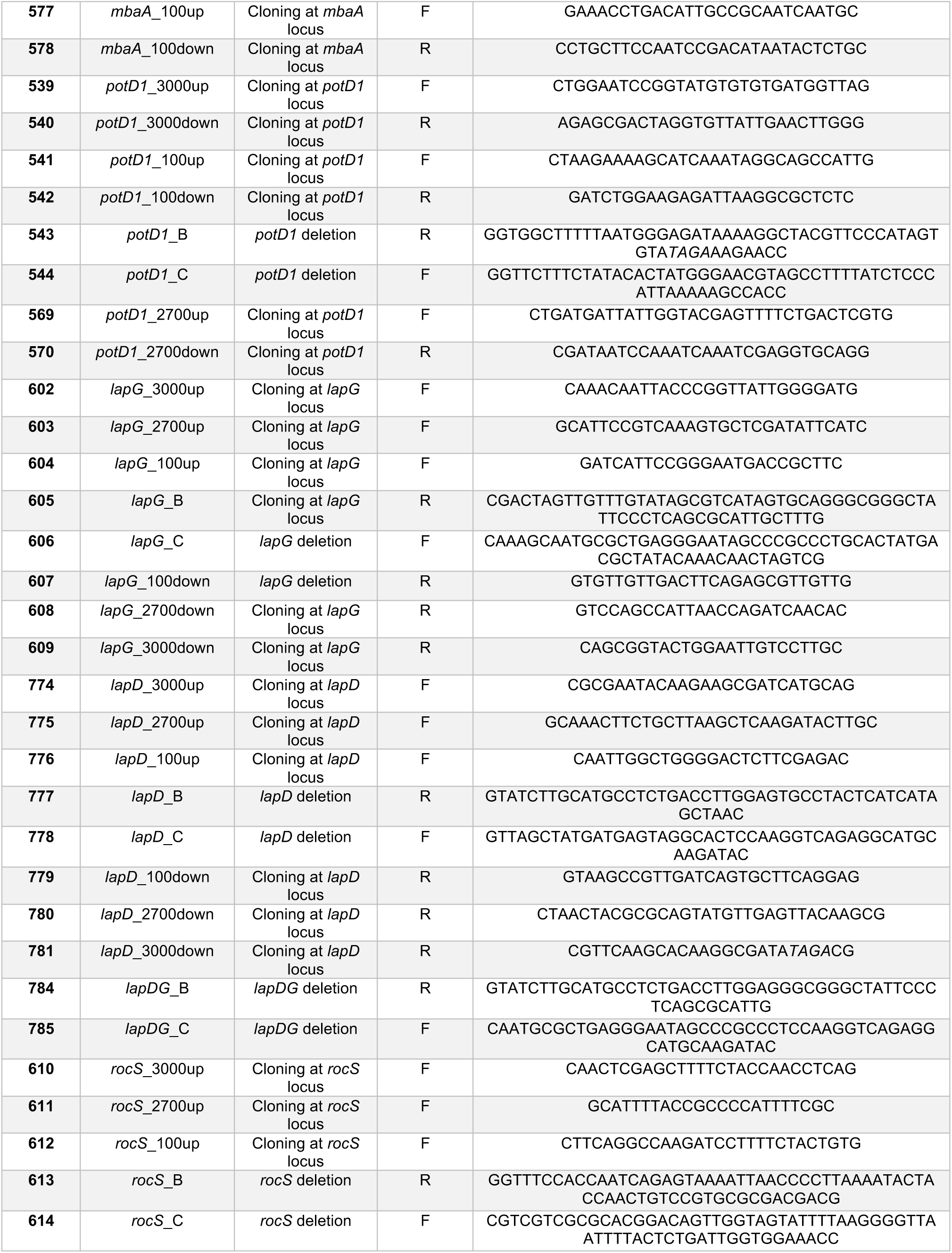
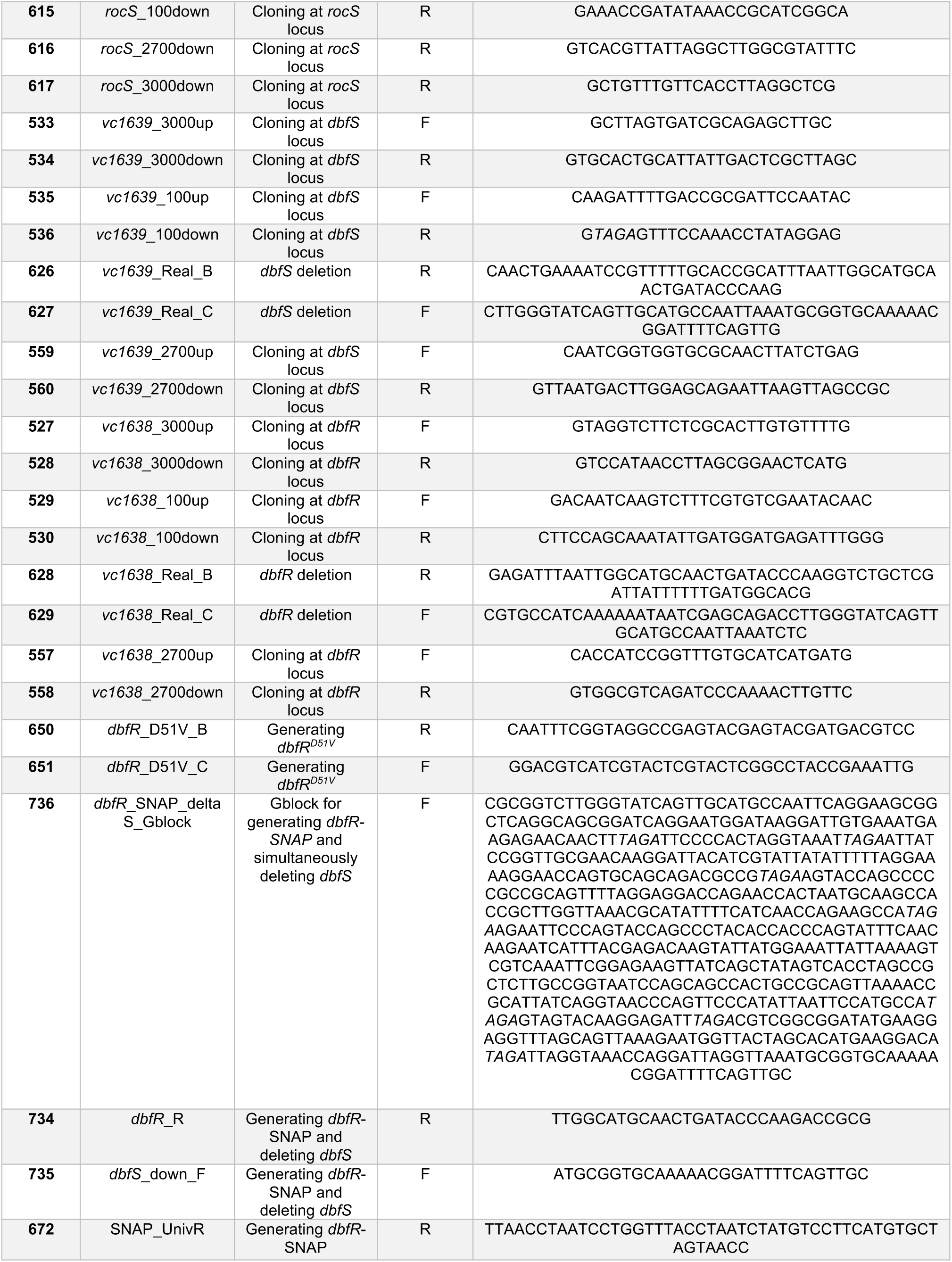
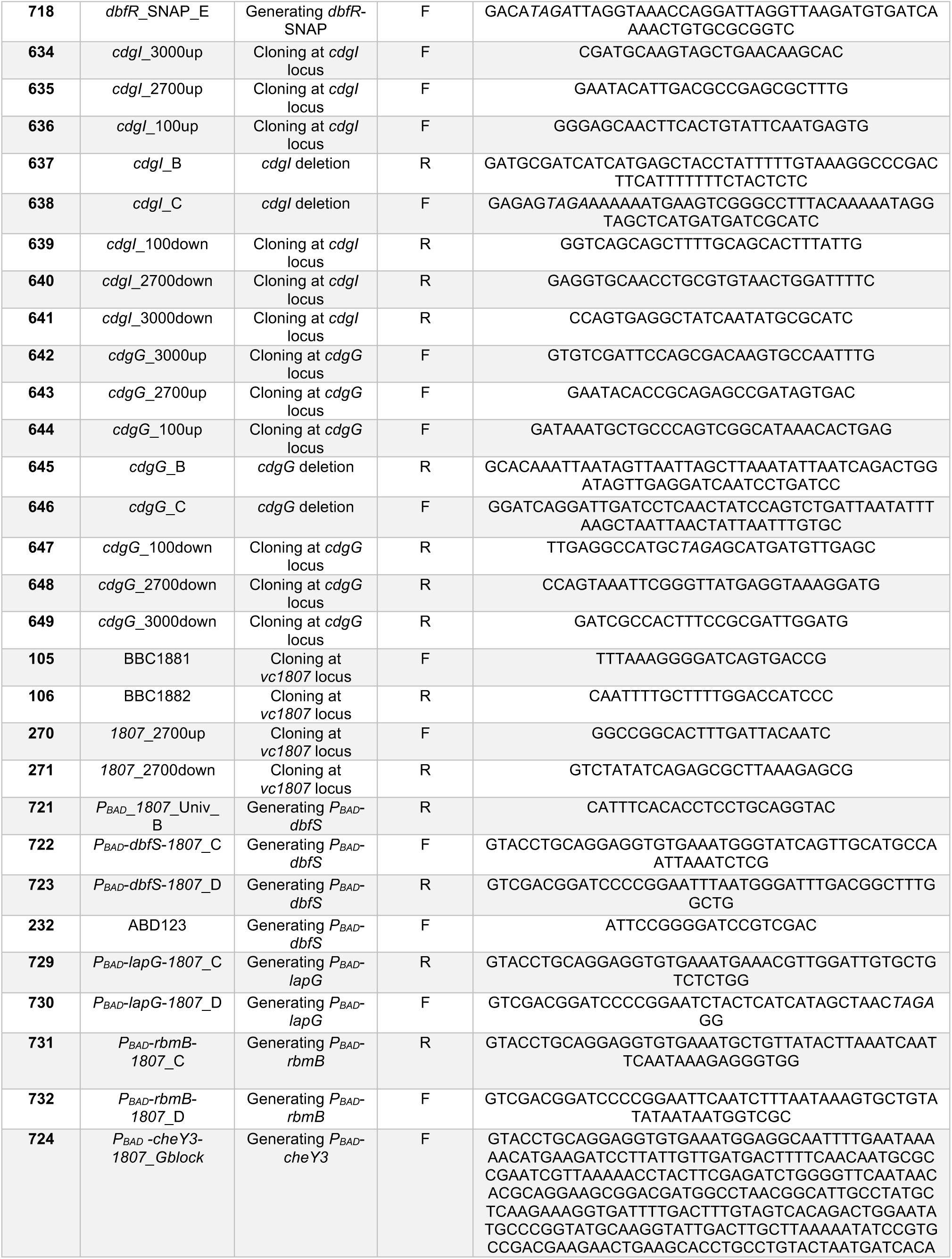
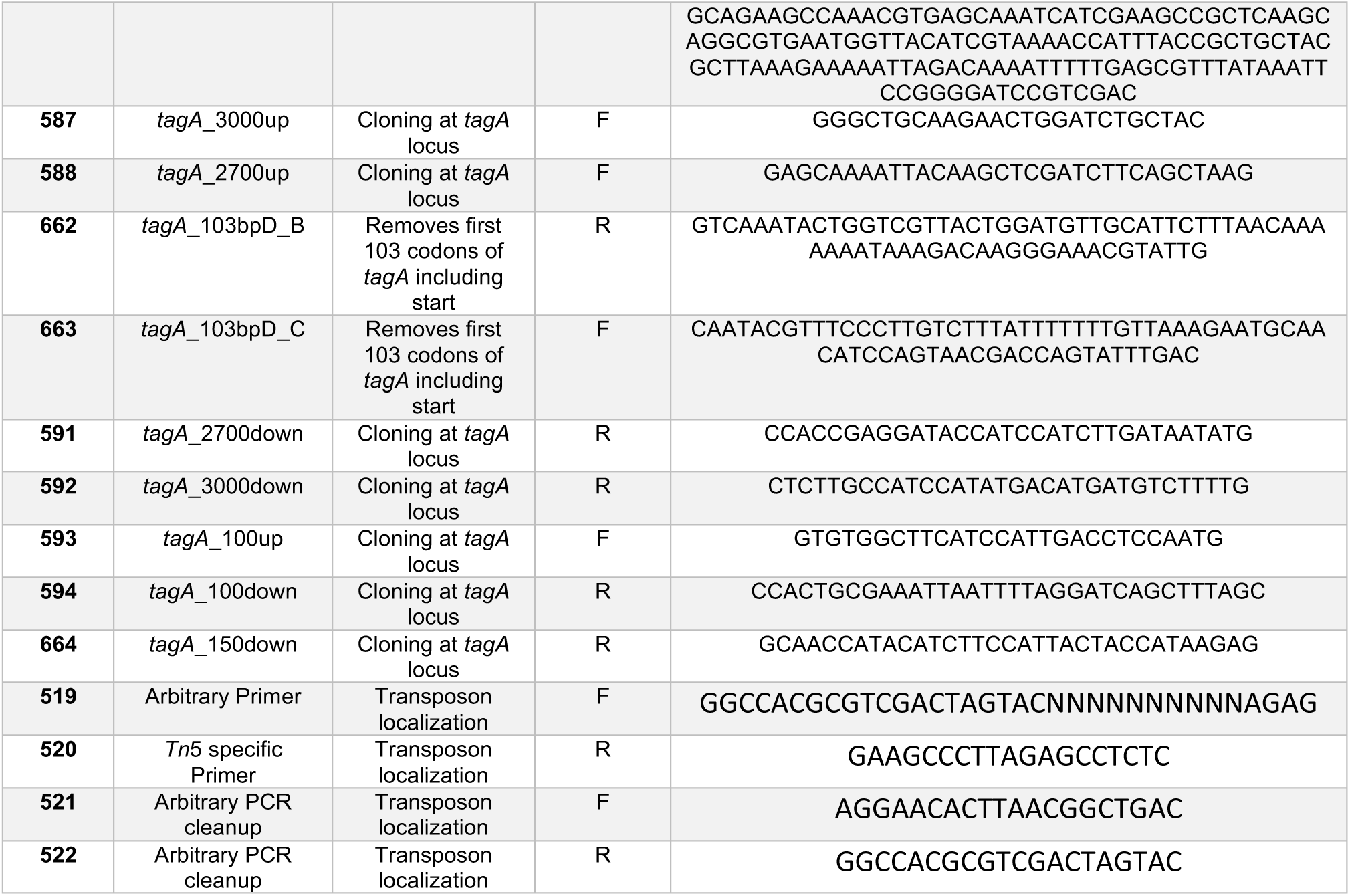
DNA oligonucleotides and gene fragments used in this study.

